# COVID-19 Neuropathology: evidence for SARS-CoV-2 invasion of Human Brainstem Nuclei

**DOI:** 10.1101/2022.06.29.498117

**Authors:** Aron Emmi, Stefania Rizzo, Luisa Barzon, Michele Sandre, Elisa Carturan, Alessandro Sinigaglia, Silvia Riccetti, Mila della Barbera, Rafael Boscolo-Berto, Patrizia Cocco, Veronica Macchi, Angelo Antonini, Monica De Gaspari, Cristina Basso, Raffaele De Caro, Andrea Porzionato

**Affiliations:** Institute of Human Anatomy, Department of Neuroscience, University of Padova, Italy; Department of Cardio-Thoracic-Vascular Sciences & Public Health, University of Padova, Italy; Department of Molecular Medicine, University of Padova, Padova, Italy; Movement Disorders Unit, Neurology Clinic, Department of Neuroscience, University of Padova; Pathology and Histopathology Unit, Ospedali Riuniti Padova Sud, Padova, Italy

## Abstract

Neurological manifestations are common in COVID-19, the disease caused by SARS-CoV-2. Despite reports of SARS-CoV-2 detection in the brain and cerebrospinal fluid of COVID-19 patients, it’s still unclear whether the virus can infect the central nervous system, and which neuropathological alterations can be ascribed to viral tropism, rather than immune-mediated mechanisms.

Here, we assess neuropathological alterations in 24 COVID-19 patients and 18 matched controls who died due to pneumonia / respiratory failure. Aside from a wide spectrum of neuropathological alterations, SARS-CoV-2-immunoreactive neurons were detected in specific brainstem nuclei of 5 COVID-19 subjects. Viral RNA was also detected by real-time RT-PCR. Quantification of reactive microglia revealed an anatomically segregated pattern of inflammation within affected brainstem regions, and was higher when compared to controls. While the results of this study support the neuroinvasive potential of SARS-CoV-2, the role of SARS-CoV-2 neurotropism in COVID-19 and its long-term sequelae require further investigation.

## Introduction

Neurological manifestations are common in coronavirus disease 19 (COVID-19), the disease caused by severe acute respiratory syndrome coronavirus-2 (SARS-CoV-2)^1–5^. Symptoms range from anosmia, ageusia, dizziness and headache, which are commonly reported by patients with mild disease, to altered mental status, neuropsychiatric disorders, stroke, and, rarely, meningitis, encephalitis, and polyneuritis, which occur in hospitalized patients with severe disease^1, 5^. Between 10 to 30% of people with SARS-CoV-2 infection experience long-term sequelae, referred as “long COVID”, including neurological manifestations such as hyposmia, hypogeusia, headaches, fatigue, sleep disorders, pain, and cognitive impairment^3^. Despite some reports of detection of SARS-CoV-2 in the brain and cerebrospinal fluid of patients with COVID-19^2–3, 6^, it is still unclear whether the virus can infect the central nervous system (CNS). In particular, it still remains to be elucidated whether neurological manifestations and neural damage are a direct consequence of viral invasion of the CNS, are due to post-infectious immune-mediated disease, or are the result of systemic disease^1, 6–11^. Studies on human neural cell cultures and brain organoids report conflicting data on SARS-CoV-2 neurotropism^12^. Overall, they suggest that SARS-CoV-2 does not infect and replicate efficiently in human neural cells, while it can replicate at high rates in choroid plexus epithelial cells^7, 13–14^. At variance, intranasal inoculation of SARS-CoV-2 in transgenic mice overexpressing human ACE2 under the K18 promoter resulted in brain invasion and widespread infection of neurons, radial glia and neuronal progenitor cells^15–16^. Other coronaviruses, such as SARS-CoV and MERS-CoV, appear to be able to infect the CNS in both humans and animal models^17^.

Data deriving from large autopsy studies in patients who died from COVID-19 suggest for the neuroinvasive potential of SARS-CoV-2 in the CNS^8–9, 17^, even though infection appears to be limited to sparse cells in the brainstem and not associated with encephalitis or other specific changes referable to the virus^8^. Conversely, other studies failed to detect SARS-CoV-2 antigens or genomic sequences in brain tissues of COVID-19 patients^11, 17–18, 37–38^. In numerous instances, neuropathological changes in the brains of COVID-19 patients were moderate and mainly represented by ischaemic lesions, astrogliosis, microglial nodules, and cytotoxic T lymphocyte infiltrates, most pronounced in the brainstem, cerebellum, and meninges^8–9, 11, 18, 21^. While diffuse to focal hypoxic / ischaemic damage was a common finding in COVID-19 patients across studies, no direct link between encountered neuropathological alterations and direct viral invasion could be established, with systemic inflammation and hypoxia playing a likely major role in mediating brain immune response^37^. Single-nucleus gene-expression profiling of frontal cortex and choroid plexus tissues from severe COVID-19 patients showed broad perturbations, with upregulation of genes involved in innate antiviral response and inflammation, microglia activation and neurodegeneration^20^, but no direct evidence of viral tropism was found; similarly, Fullard et al.^38^ were unable to detect viral transcripts and S proteins in different brain regions of COVID-19 subjects. Deep spatial profiling of the local immune response in COVID-19 brains through imaging mass spectrometry revealed significant immune activation in the CNS with pronounced neuropathological changes (astrocytosis, axonal damage, and blood-brain-barrier leakage) and detected viral antigen in ACE2-positive cells enriched in the vascular compartment^18^. According to the study, the presence of viral antigen was linked to vascular proximity and ACE2 expression, while also being correlated to the perivascular immune activation patterns of CD8 and CD4 T cells and myeloid- and microglial-cell subsets^18^, indicating a fundamental role of the vascular and perivascular compartment, as well as Blood-brain-barrier impairment, in mediating COVID-19 specific neuropathological changes. As evidenced by the above case series, and considering the different case reports available^22–24^, SARS-CoV-2 infection of CNS seems to be limited to isolated cells within the perivascular compartment of the brainstem and olfactory bulb, and have been reported in a subset of cases in the various autopsy series, while widespread neuropathological sequelae (such as astrogliosis, microgliosis, lymphocyte infiltration, microvascular injury, fibrinogen leakage) have been documented in most examined specimens. The possibility of direct viral invasion, and eventual associated long-term sequelae of infection, remain to be investigated.

In the present study, we assess the neuropathological changes of 24 patients who died following a diagnosis of SARS-CoV-2 infection in Italy during the COVID-19 pandemic (from March 2020 to May 2021) and 18 age-matched controls with comparable medical conditions who died mainly due to pneumonia and / or respiratory failure.

## Study design and Materials

Hospitalized patients who died following a diagnosis of SARS-CoV-2 infection in the Veneto Region, Italy, during the peak incidence of COVID-19 (from March 2020 to May 2021) were autopsied according to established COVID-19 infection security protocols. Inclusion criteria for the study were: a) diagnosis of SARS-CoV-2 infection confirmed by molecular testing of rhino-pharyngeal swabs and b) high-quality brain tissue samples available for histopathological and immunohistochemical analysis. Tissue quality was determined by Post-Mortem Interval (PMI) ≤5 days, absence of tissue maceration, fixation time ≤ 3 weeks and adequate formalin penetration within the tissue. A total of 24 COVID-19 patients were included in the study.

18 age- and sex-matched subjects with comparable general medical conditions, predating the COVID-19 pandemic in Italy, were included as controls.

## Methods

### Clinical information

Available clinical data for COVID-19 subjects and controls were examined, including ante-mortem medical history, neurological and neuroradiological findings, hospitalization time, ICU and oxygen therapy status, and prescribed medication.

However, as most subjects died during the sanitary emergency of the first wave of the COVID-19 pandemic in Italy, ante-mortem clinical data were at times limited, especially when concerning post-hospitalization neurological status. This represents one of the main limitations of our study, determining significant constrains to the association between ante-mortem neurological findings and encountered neuropathological alterations, which is often not unequivocal.

### Sampling and fixation procedures

Sampled brains were immersion fixed in 4% phosphate-buffered formalin solution following autopsy (mean PMI: 3 days; Range 0-5 days; average fixation time: 2-3 weeks) and subsequently sectioned for histopathological and immunohistochemical analysis. Samples of the cerebral cortex, basal ganglia, hippocampus, cerebellar cortex, deep cerebellar nuclei, choroid plexuses and meninges were obtained, while the brainstem was isolated at the level of the rostral extremity of the midbrain and extensively sampled in its whole cranio-caudal extent. The 12 cranial nerves, where available, including the olfactory bulb, tract and bifurcation, were also sampled. To preserve antigen quality, a slow dehydration and clearing protocol was performed prior to paraffin embedding (24h mean tissue processing time).

### Histochemical and immunoperoxidase staining

Haematoxylin and Eosin staining was employed for routine histopathological evaluation. Immunoperoxidase staining was performed on a Dako EnVision Autostainer (Dako Denmark A/S, Glostrup, Denmark) according to manufacturer recommendations. Antibodies for CD3 (Polyclonal Rabbit Anti-Human, Citrate Buffer HIER, dilution 1:200, Dako Omnis, Code Number: GA503), CD20 (Monoclonal Mouse Anti-Human, Citrate Buffer HIER, dilution 1:200 Clone KP1, Dako Omnis, Code Number: M0814) and CD68 (Monoclonal Mouse Anti-Human, EDTA Buffer HIER, IHC dilution 1:5000, IF dilution 1:500, Clone L26, Dako Omnis, Code Number: M0756) were employed to characterize lympho-monocytic infiltrations. Microglial Activation was assessed using both CD68 (as above), HLA-DR Antibody (Monoclonal Rabbit Anti-Human, Citrate Buffer HIER, dilution 1:50 Clone: LN-3, Invitrogen, Thermo Fisher Scientific, Waltham, MA, USA), TMEM119 (Rabbit Anti-Human, Citrate Buffer HIER, dilution 1:250, Abcam, Code Number: ab185333), while microglial proliferation was assessed using anti-Ki-67 immunohistochemistry (Mouse Anti-Human, EDTA Buffer HIER, dilution 1:200, Spring Bioscience, Code number: M3060). Anti-GFAP immunohistochemistry (Polyclonal Rabbit Anti-Human, Proteinase K enzymatic antigen retrieval, dilution 1:1000, DAKO Omnis, Code Number: GA524) was employed to assess reactive astrogliosis. Anti-CD61 immunohistochemistry (Monoclonal Mouse Anti-Human, Citrate Buffer HIER, dilution 1:75, Clone Y2/51, Dako Omnis, Code Number: M0753) was also employed to evaluate the presence of platelet-enriched microthrombi. Anti-SARS-CoV-2 nucleocapsid (Rabbit Anti-Human, Citrate Buffer HIER, dilution 1:7000, Sino Biologicals, 40143-R001) and -Spike Subunit 1 Antibody (Monoclonal Rabbit Anti-Human, Citrate Buffer HIER, dilution 1:100, Clone 007, Sino Biological, Code Number: 40150-R007) immunostainings were employed to evaluate viral antigens within the tissue. The expression of ACE2 Receptor protein (Rabbit Anti-Human Polyclonal, Citrate Buffer HIER, dilution 1:2000, Abcam, Code Number: ab15348) and TMPRSS-2 protein (Rabbit Anti-Human Monoclonal, Citrate Buffer HIER, dilution 1:2500, Abcam, Code Number: ab242384) was assessed within the brainstem and cerebellum, and in all sections with positive findings for viral proteins. Anti-nucleocapsid and anti-spike antibodies were validated through SARS-CoV-2 infected Vero E6 cells and autopsy-derived lung tissue from SARS-CoV-2 infected patients as positive controls; non-infected cells and lung sections deriving from autopsy cases predating COVID-19 pandemic (2017) were used as negative controls (Supplementary Figure 1). Peroxidase reactions were repeated at least three times to ensure reaction consistency.

### Immunofluorescent staining and confocal microscopy

Fluorescent immunohistochemistry was performed manually. Antigen retrieval was performed on de-paraffinized tissue sections using Dako EnVision PTLink station according to manufacturer recommendations. Following antigen retrieval, autofluorescence was quenched with a 50 mM NH4Cl solution for 10 minutes. Sections were treated with permeabilization and blocking solution (15% vol/vol Goat Serum, 2% wt/vol BSA, 0.25% wt/vol gelatin, 0.2% wt/vol glycine in PBS) containing 0.5% Triton X-100 for 90 minutes before primary antibody incubation. The following antibodies were employed: CD68 (#M0756; 1:500); TMEM119 (#ab185333; 1:200); Ki-67 (#M3060; 1:200); β-III Tubulin (#T8578; 1:300); Tyrosine Hydroxylase (#T2928; 1:6000); SARS-CoV-2 Nucleocapsid Protein (#40143-R001; 1:3000); SARS-CoV-2 Spike Subunit 1 Protein (#40150-R007; 1:100); ACE2 Receptor Protein (#ab15348; 1:500) and TMPRSS-2 (#ab242384; 1:1000). Primary antibodies were diluted in blocking solution and incubated at 4°C overnight. Alexa-Fluor plus 488 Goat anti-Mouse secondary antibody (Code number: A32723) and Alexa-Fluor plus 568 anti-Rabbit secondary antibody (Code number: A-11011) were diluted 1:200 in blocking solution as above and incubated for 60 minutes at room temperature. To further avoid background signal and tissue autofluorescence, slides were incubated for 10 minutes in 0.5% Sudan Black B solution in 70% ethanol at room temperature and abundantly washed with PBS, followed by Hoechst 33258 nuclear staining (Invitrogen, dilution: 1:10000 in PBS) for 10 minutes. Slides were mounted and coverslipped with Mowiol solution (prepared with Mowiol 4-88 reagent, MerckMillipore, Code number: 475904-100GM). Confocal immunofluorescence z-stack images were acquired on a Leica SP5 Laser Scanning Confocal Microscope using a HC PL FLUOTAR 20x/0.50 Dry or HCX PL APO lambda blue 40X/1.40 Oil objectives. Images were acquired at a 16-bit intensity resolution over 2048 × 2048 pixels. Z-stacks images were converted into digital maximum intensity z-projections, processed, and analyzed using ImageJ software.

### RT-PCR analyses

Viral RNA analysis was performed on 20µm thick paraffin-embedded sections collected in sterile 2ml Eppendorf vials; disposable microtome blades and tongs were changed for each section to reduce contamination risk. Real-time RT-PCR analyses were performed to detect SARS-CoV-2 genome sequences. Briefly, total RNA was purified from selected material using a RecoverAll™ Total Nucleic Acid Isolation kit (Thermo Fisher Scientific) following the manufacturer’s instructions. One-step real-time RT-PCR assays targeting SARS-CoV-2 nucleocapsid (N) coding region and subgenomic RNA were run on ABI 7900HT Sequence Detection Systems (Thermo Fisher Scientific), as previously reported^25^.

### Histopathological and morphometrical evaluation

Slides were examined by three independent histopathologists and morphologists blind to patient clinical findings and COVID-19 status. Disagreements were resolved by consensus. The degree of brainstem hypoxic / ischaemic damage, astrogliosis and microgliosis were classified using a four-tiered semi-quantitative approach for each evaluated section, while microglial density and activation was assessed by the means of digitally-assisted immunoreactivity quantification by three independent evaluators.

### Quantification of Activated Microglia

The degree of microgliosis was assessed through a digitally-assisted quantification approach at the level of the medulla, pons and mesencephalon. For each subject, standard sections passing through the area postrema (medulla), locus coeruleus (pons) and decussation of the superior cerebellar peduncles or red nucleus (midbrain) underwent TMEM119 immunoperoxidase staining and TMEM119 / CD68 double fluorescent immunohistochemistry. TMEM119+ structures with visible nucleus and microglial-compatible morphology were classified as microglial cells, while TMEM119-/ CD68+ elements with compatible morphology were classified as monocyte/macrophages. Ramifications and cell processes without a visible nucleus were excluded from our analysis in order to avoid overestimation of cell densities by including neighboring structures belonging to adjacent sections. Morphometrical evaluation occurred within six counting fields (fields of view, FOV) spanning across the dorsal-to-ventral axis of the sections; FOV boundaries and anatomical landmarks are summarized in Supplementary Table 1 for each level of sectioning. The number of immunoreactivities per mm^2^ was calculated for each counting field and assigned to one anatomical compartment (i.e. tegmentum, tectum and basis), based on their topography according to Mai and Paxinos^26^. Comparisons and statistical evaluations were conducted per individual counting field, anatomical compartment and level of section (medulla, pons, midbrain). To assess the degree of lysosomal-activity as a marker for microglia phagocytic activity, CD68 immunoreactive area (expressed as percentage of CD68+ immunoreactive area within a counting field, or A%) for 5 randomly selected counting fields at each level of sectioning was computed through particle analysis of the green fluorescent channel on ImageJ software.

### Statistical Analyses

Statistical analyses and visualizations were performed using GraphPad Prism 9. Differences in microglial densities (microglia / mm^2^) within subgroups of the COVID-19 cohort in figures 4A, 5A and 6A were analyzed by t tests with Welch’s correction. Microglial density between individual counting fields (FOVs) in COVID-19 subjects (Figures 4D, 5D, and 6D), as well as differences between anatomical compartments in COVID-19 subgroups (Figures 4B and 5B) and in COVID-19 versus controls (Figures 2E, 4C, 5B, 6E) were determined by Welch one-way ANOVA tests corrected for Dunnett’s multiple comparisons. Correlation matrices in Figures 4E and 7G were computed as Spearman’s rho for continuous variables and as point-biserial correlations for nominal – continuous variables. Spearman’s rho and linear regression was performed in figure 2F. Further statistical details for each plot can be found in the corresponding figure legend. Throughout the manuscript * indicates p < 0.05, ** p < 0.01, *** p < 0.001 and **** p < 0.0001.

**Figure 1.**
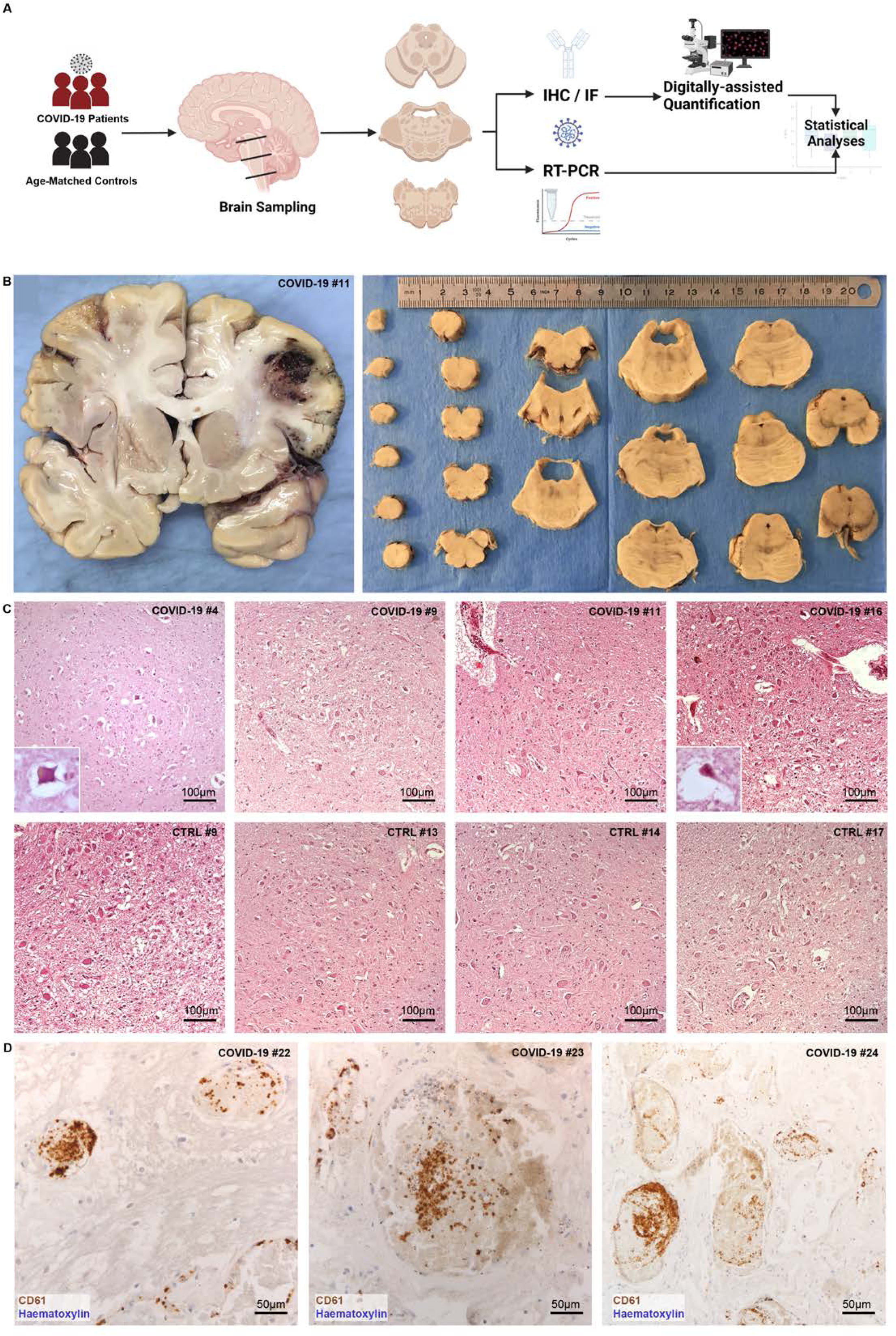
**A)** Study Workflow. Brain sections of multiple sites were sampled from 24 COVID-19 patients and 18 age- and sex-matched controls who died due to pneumonia and / or respiratory failure. **B)** Left, coronal brain section of Subject #11 revealing extensive hemorrhagic injury in the territory of the middle cerebral artery. Right, sampling procedure of the brainstem through axial sections passing perpendicularly to the floor of the fourth ventricle. **C)** Haematoxylin and eosin photomicrographs of the Dorsal Motor Nucleus of the Vagus in the medulla oblongata displaying various degrees of hypoxic / ischaemic damage in COVID-19 subjects (upper row) and controls (lower row) **D)** Platelet microthrombi at the level of the pons and cerebral cortex, CD61 immunohistochemistry.

**Figure 2.**
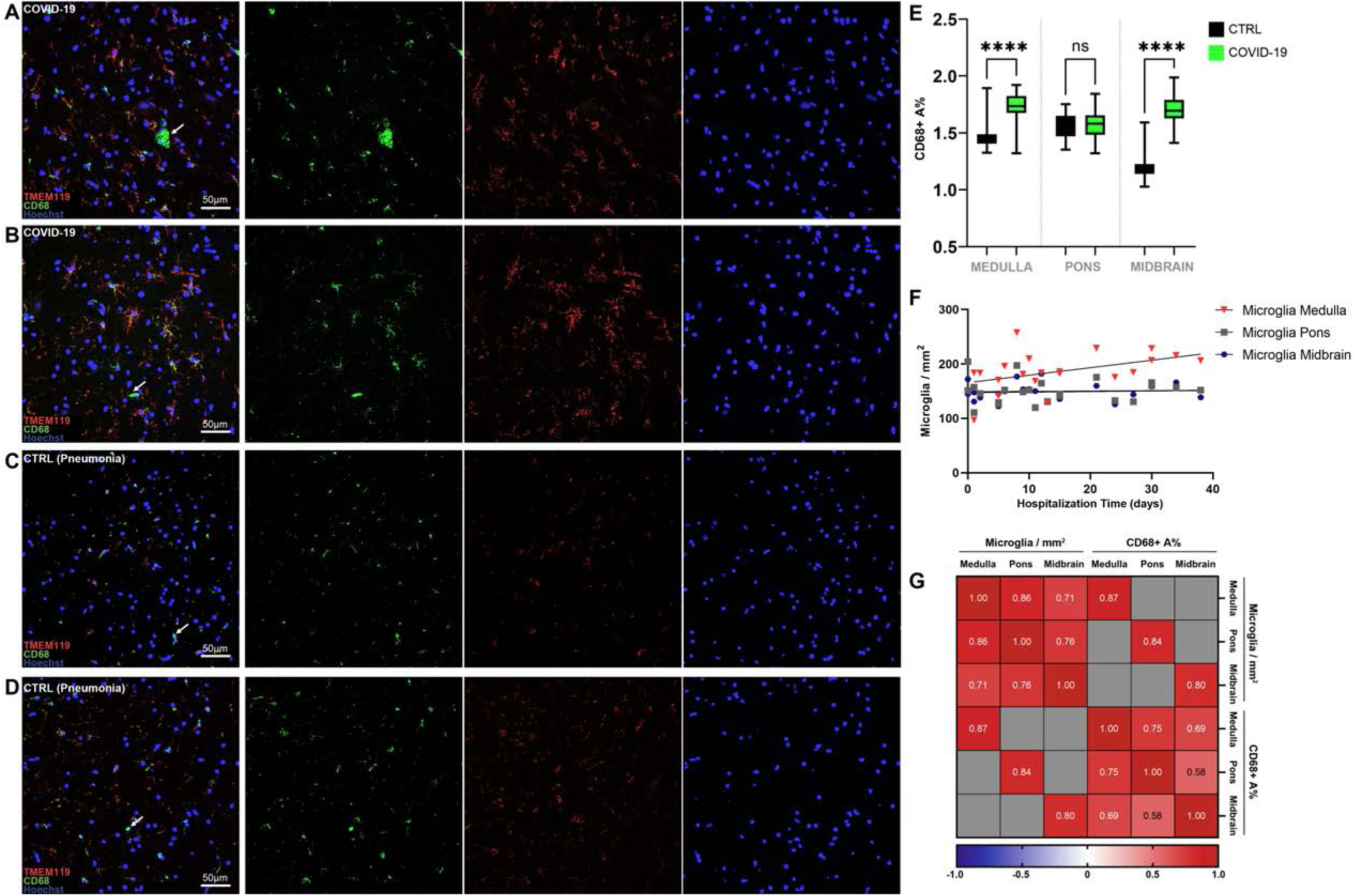
Double label TMEM119 (microglia marker, red) / CD68 (lysosomal activity marker, green) fluorescent immunohistochemistry in COVID-19 and control subjects. **A-B)** in COVID-19, microglial cells present a distinctly activated phenotype whilst maintaining homeostatic microglial marker TMEM119 (Red) and displaying increased lysosomal activity (CD68, green). Arrows indicate CD68+ / TMEM119-monocyte/macrophage in the parenchyma. **C-D)** In control subjects, TMEM119 marks both the soma and sparse ramifications of resident microglia, suggesting less prominent activation without significant marker downregulation. CD68 immunoreactivity (green) is also present, but not as distributed as in COVID-19. **E)** Welch one-way ANOVA of CD68+ A% in COVID-19 and controls reveals statistically significant differences between the two groups at the level of the medulla (p<0.0001) and midbrain (p<0.0001), but not the pons. **F)** Spearman correlation between microglial densities across brainstem levels and hospitalization time reveals a statistically significant positive correlation between medullary microgliosis and hospitalization time (r= 0.44; p=0.044). **G)** Correlation matrix between microglial densities and CD68+ A% across brainstem levels.

**Figure 3.**
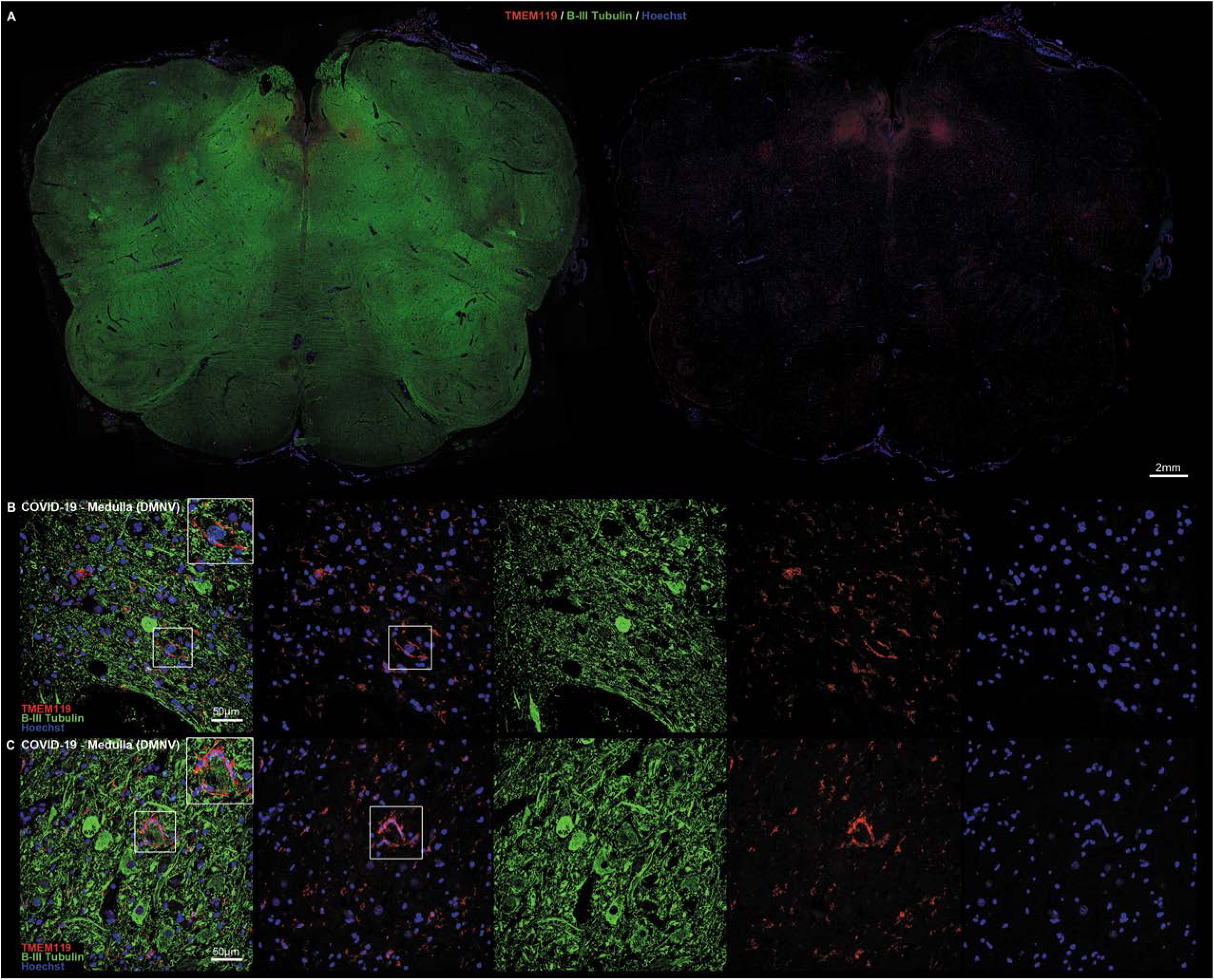
**A)** Low-magnification perspective of the medulla oblongata in COVID-19, displaying a ventral-to-dorsal increasing gradient of microglial densities (TMEM119, red; Beta-III Tubulin, green) **B-C)** Double label TMEM119 (microglial marker, red) and Beta-III Tubulin (neuronal marker, green) immunofluorescent staining at the level of the medulla oblongata. Insets display neuronophagia at the level of the dorsal motor nucleus of the vagus in two COVID-19 subjects.

**Figure 4.**
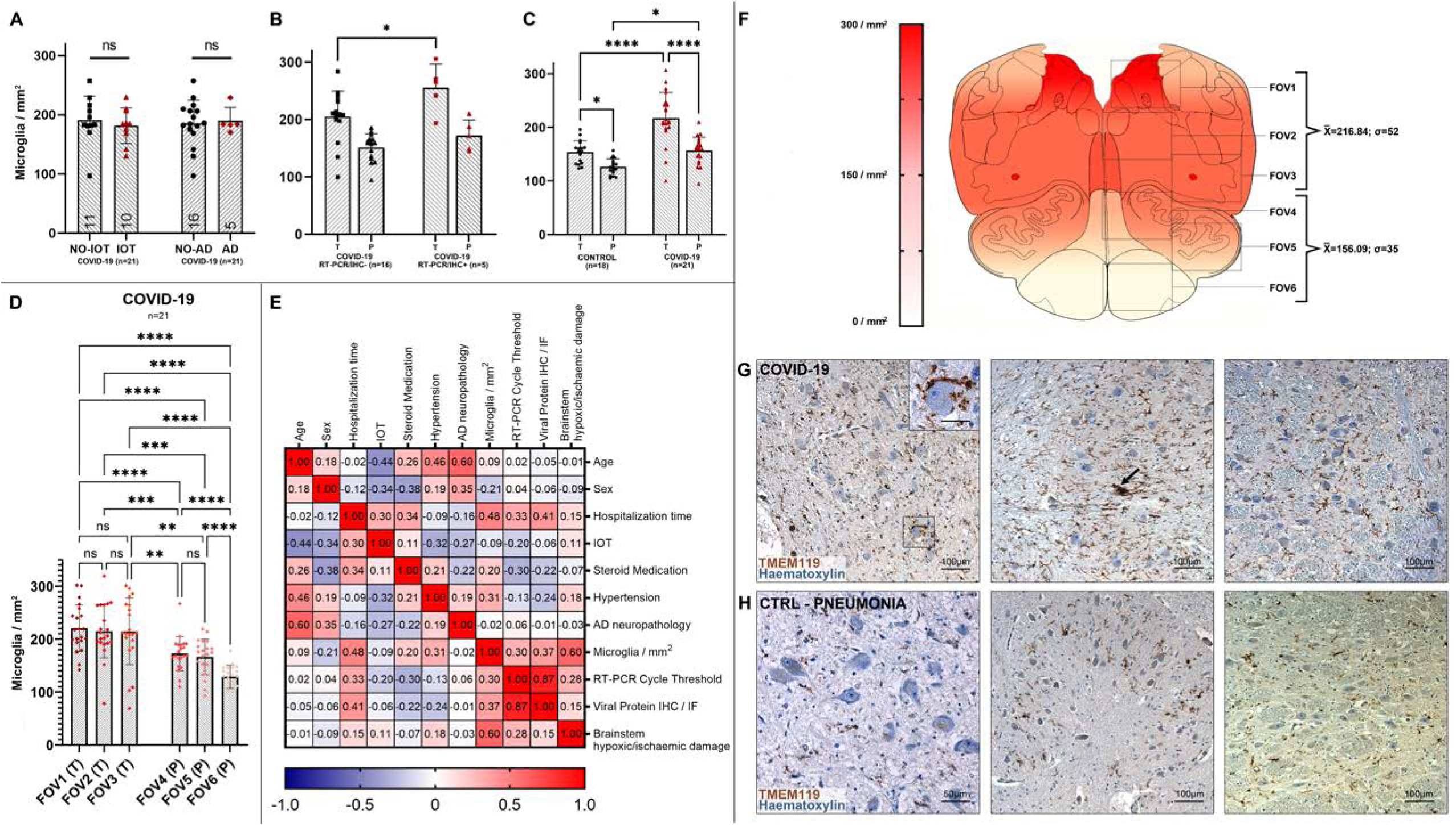
**A)** Welch corrected T-test plot of microglial densities (microglia / mm^2^) in the medulla oblongata of COVID-19 subjects treated (n=10, red) and not treated (n=11, black) with intensive oxygen therapy (p>0.05), and of COVID-19 subjects with (n=5, red) and without (n=16, black) Alzheimer’s Disease neuropathological changes (p>0.05). **B)** Welch one-way ANOVA between anatomical compartments (T, tegmentum; P, pes) between COVID-19 subjects with (n=5, red) and without (n=16) viral tropism (RT-PCR/IHC+ versus RT-PCR/IHC-) reveals statistically significant differences at the level of the medullary tegmentum (p=0.017). **C)** Welch one-way ANOVA between anatomical compartments (T, tegmentum; P, pes) between COVID-19 subjects (n=21, red) and controls (n=18, black) reveals statistically significant differences both at the level of the medullary tegmentum (p<0.0001) and pes (p=0.017). **D)** Welch one-way ANOVA of microglial densities per counting fields (FOV) reveals statistically significant differences between FOVs of the Tegmentum (T; FOV1-3) when compared to FOVs of the Pes (P; FOV4-6). **E)** Correlation heatmap between COVID-19 subject clinical data and neuropathological findings. **F)** Anatomical heatmap of activated microglia within the medulla oblongata in COVID-19. **G-H)** TMEM119 immunoperoxidase staining of comparable regions of the medulla oblongata in COVID-19 subjects (above) and controls (below). Inset: neuronophagia in the dorsal motor nucleus of the vagus. Arrow: microglial nodule.

**Figure 5.**
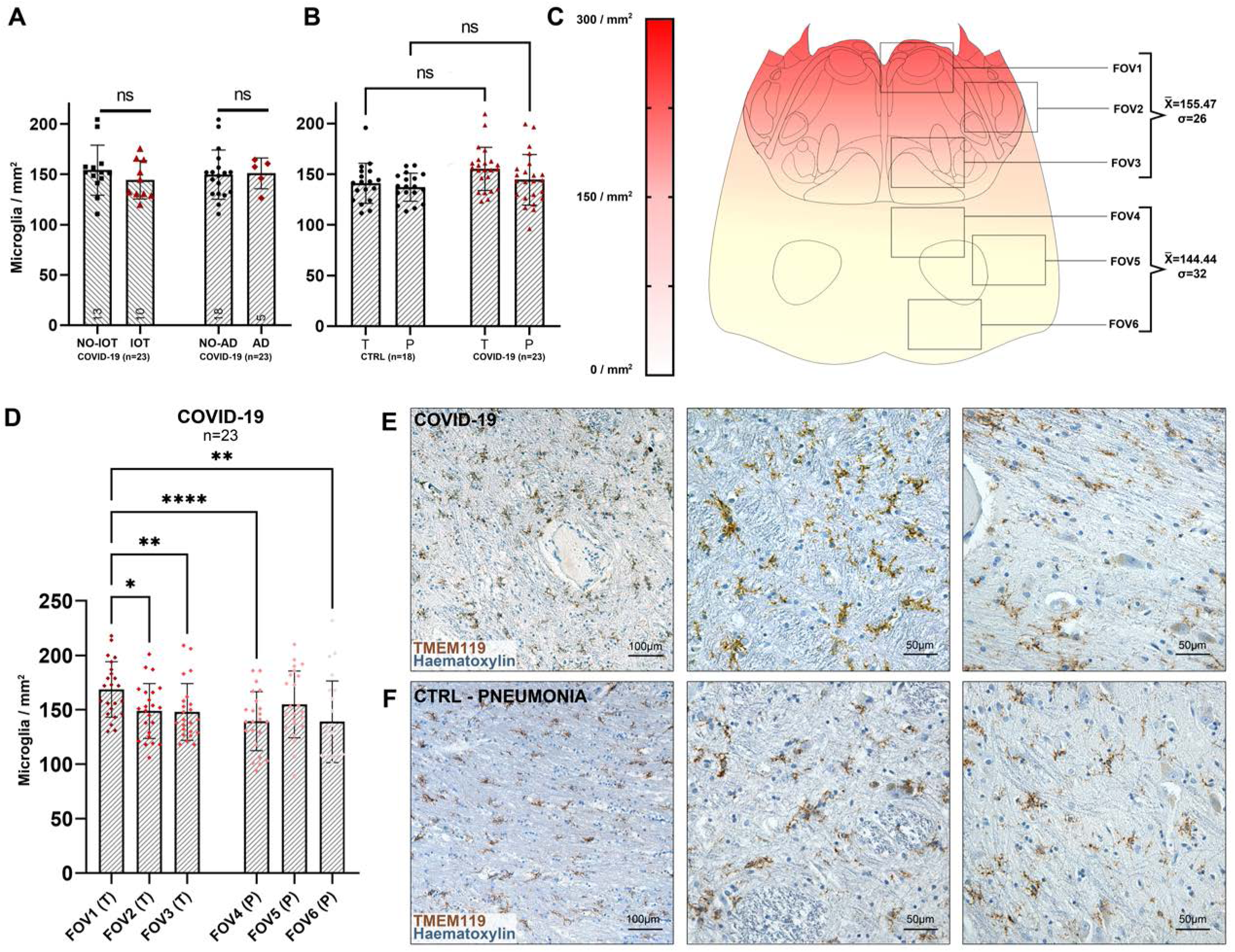
**A)** Welch corrected T-test plot of microglial densities (microglia / mm^2^) in the pons of COVID-19 subjects with (n=5, red) and without (n=16, black) Alzheimer’s Disease neuropathological changes (p>0.05), and of COVID-19 subjects treated (n=10, red) and not treated (n=13, black) with intensive oxygen therapy (p>0.05). **B)** Welch one-way ANOVA between anatomical compartments (T, tegmentum; P, pes) between COVID-19 subjects (n=23, red) and controls (n=18, black) reveals no statistically significant differences either at the level of the medullary tegmentum (p=0.55) and pes (p=0.98). **C)** Anatomical heatmap of activated microglia within the pons in COVID-19. **D)** Welch one-way ANOVA of microglial densities per counting fields (FOV) reveals statistically significant differences between FOV1 (dorsal pons, including locus coeruleus) with other pontine counting fields. **E-F)** TMEM119 immunoperoxidase staining of comparable regions of the pons in COVID-19 subjects (above) and controls (below).

## Results

### The main cause of death in COVID-19 subjects was diffuse alveolar damage

Twenty-four COVID-19 patients were included in our study. In all patients, SARS-CoV-2 RNA was detected by molecular testing in rhino-pharyngeal swabs. Eleven were females, while 13 were males. The mean age of the included subjects was 73±13.7 years. Most included subjects were affected by preexisting chronic medical conditions. Eleven patients (7 female, 4 male) were affected by neurological or neurodegenerative disease prior to SARS-CoV-2 infection. Twenty-three patients were hospitalized prior to death. Patients were hospitalized for 14.5±11.3 days and died 1 to 34 days following admission. Eleven subjects were admitted to the ICU during hospitalization and received intensive oxygen therapy (IOT) (i.e. the administration of supplemental oxygen via nasal cannulae, face masks, or tracheal intubation). Fifteen subjects received antithrombotic therapy during hospitalization and were treated with corticosteroid medication. The available clinical data for our cohort is reported in Table 1.

**Table 1.**
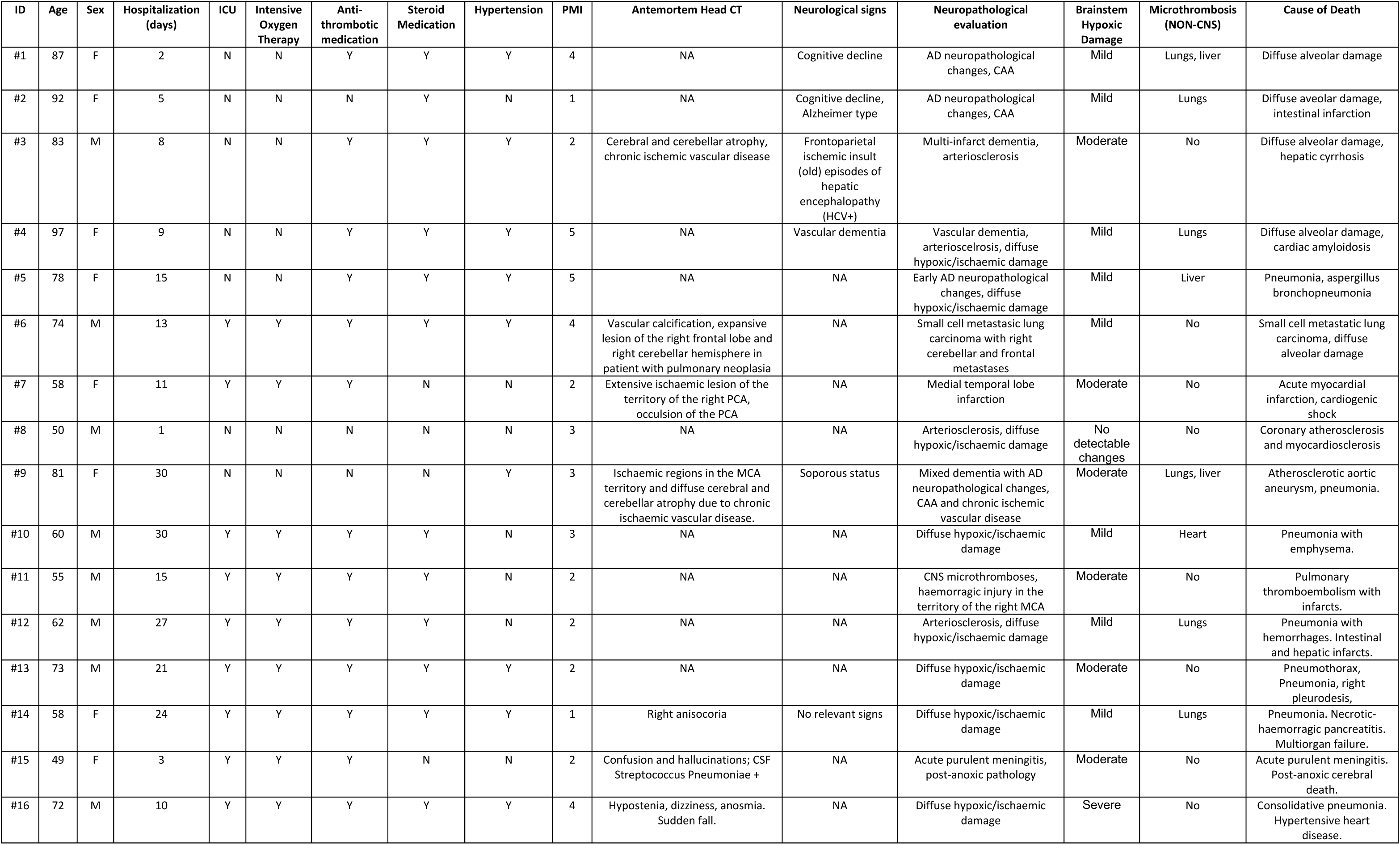
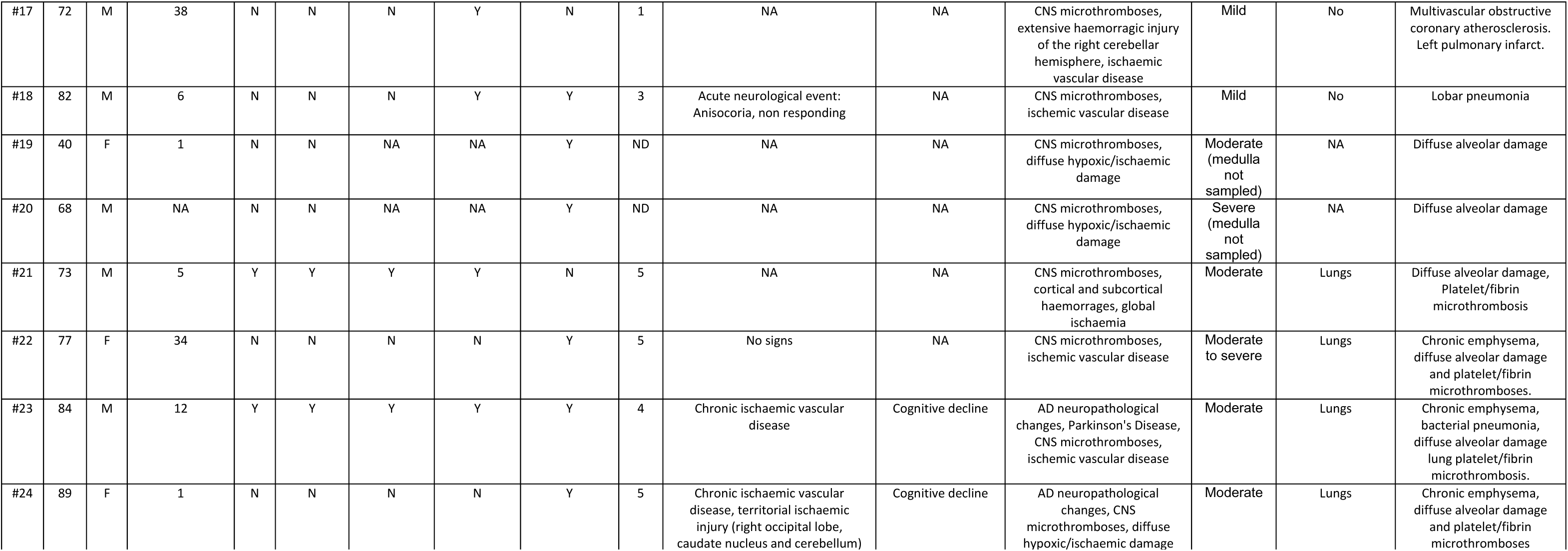
Clinical data of the COVID-19 Group.

### The main cause of death of the control cohort was respiratory failure and pneumonia, aside from other relevant comorbidities

Eighteen age- and sex-matched subjects with comparable ante-mortem medical conditions were included as controls. All patients were negative for SARS-CoV-2 infection or died prior to the COVID-19 pandemic in Italy. Eight were female, while 10 were male. The mean age of included controls was 72±12 years. The mean hospitalization time was 20±15.6 days. Thirteen patients died due to pneumonia, while the remaining subjects died due to respiratory insufficiency, multiorgan failure or ischaemic heart disease. One patient died due to septic shock. Five patients had a clinical diagnosis of cognitive decline. The available clinical data for the control group are reported in Table 2.

**Table 2.** Clinical data of the COVID-19 Group.

| ID | Age | Sex | Hospitalization (days) | Hypertension | PMI | Antemortem Head CT | Neurological signs | Neuropathological evaluation | Brainstem Hypoxic damage | Microthrombosis (NON-CNS) | Cause of Death |
| --- | --- | --- | --- | --- | --- | --- | --- | --- | --- | --- | --- |
| #1 | 83 | M | 10 | Y | 5 | Cerebral atrophy, Chronic ischaemic vascular disease | Cognitive decline | Mixed dementia with AD neuropathological changes ad chronic ischaemic vascular disease | Moderate | No | Pneumonia, respiratory insufficiency, ischaemic heart disease |
| #2 | 74 | M | 2 | Y | 4 | Vascular calcification, ischaemic heart disease | NA | Chronic ischaemic vascular disease | Mild | No | Ischaemic heart disease. |
| #3 | 40 | M | 1 | N | 4 | No signs | No signs | No detectable microscopical changes | No detectable microscopical changes | No | Haemorrhagic Shock |
| #4 | 79 | F | 31 | Y | 3 | Cerebral atrophy, Chronic ischaemic vascular disease | Cognitive decline, Alzheimer type | AD neuropathological changes, CAA, ischemic vascular disease | Mild | No | Pentalobar pneumonia, respiratory insufficiency |
| #5 | 62 | M | 22 | Y | 5 | NA | NA | Diffuse hypoxic/ischaemic damage | Mild | No | Pneumonia, respiratory insufficiency |
| #6 | 76 | M | 15 | Y | 6 | NA | NA | Ischaemic vascular disease, diffuse hypoxic/ischaemic damage | Mild | No | Pneumonia, chronic ischaemic vascular disease |
| #7 | 75 | M | 12 | Y | 4 | NA | NA | Diffuse hypoxic/ischaemic damage | Mild | No | Pneumonia, ischaemic heart disease |
| #8 | 78 | F | 8 | Y | 3 | No | NA | Diffuse hypoxic/ischaemic damage | Mild | No | Pneumonia, ischaemic heart disease |
| #9 | 71 | F | 40 | Y | 3 | NA | NA | Diffuse hypoxic/ischaemic damage | Moderate to severe | No | Acute respiratory failure, septic shock, peritonitis |
| #10 | 46 | F | 15 | N | 4 | No signs | No signs | Diffuse hypoxic/ischaemic damage | No detectable microscopical changes | No | Respiratory insufficiency, multiorgan failure, cervical neoplasia |
| #11 | 75 | F | 20 | Y | 5 | Chronic Ischaemic Vascular disease, Cerebral atrophy | NA | Vascular dementia, ischaemic vascular disease, diffuse hypoxic/ischaemic damage | Moderate | No | Pneumonia, acute respiratory failure, candidosis |
| #12 | 80 | F | 8 | Y | 4 | Cerebral atrophy | Cognitive decline, Alzheimer type | AD neuropathological changes, diffuse hypoxic/ischaemic damage | Moderate | No | Pentalobar pneumonia, respiratory insufficiency |
| #13 | 81 | F | NA | Y | 3 | NA | NA | Diffuse hypoxic/ischaemic damage | Mild | No | Pneumonia, acute respiratory failure |
| #14 | 63 | F | NA | N | 3 | No signs | No signs | Diffuse hypoxic/ischaemic damage | Mild | No | Ischaemic heart disease |
| #15 | 70 | M | 38 | Y | 5 | Chronic ischaemic vascular disease | NA | Ischaemic vascular disease, diffuse hypoxic/ischaemic damage | Moderate | No | Bilateral pneumonia, respiratory insufficiency. |
| #16 | 81 | M | 10 | Y | 4 | Cerebral atrophy | Cognitive decline, Alzheimer type | Mixed AD neuropathological changes and Lewy Body pathology, diffuse hypoxic/ischaemic damage | Moderate | No | Pneumonia, respiratory insufficiency |
| #17 | 75 | M | 58 | Y | 6 | Cerebral atrophy | NA | Vascular dementia, ischaemic vascular disease, diffuse hypoxic/ischaemic damage | Moderate | No | Pneumonia, respiratory insufficiency. |
| #18 | 87 | M | 30 | Y | 6 | Cerebral atrophy | Cognitive decline | AD neuropathological changes, diffuse hypoxic/ischaemic damage | Moderate | No | Pneumonia, multiorgan failure. |

## Neuropathological examination

### A wide spectrum of neuropathological alterations was detected in both COVID-19 and control subjects

The brains of 20 COVID-19 subjects displayed gross macroscopic abnormalities including mild-to-moderate generalized cerebral atrophy (N=9), diffuse cerebral edema (N=9) and chronic territorial ischaemic injury (N=6). Histopathological evaluation revealed diffuse hypoxic / ischaemic damage as a common finding in the COVID-19 cohort, with most subjects presenting mild-to-moderate diffuse hypoxic / ischaemic damage of the cerebral hemispheres and brainstem (Figure 1C), quantified according a four-tiered semi-quantitative scale (reported in Table 1). Furthermore, acute ischaemic injuries were evident in 5 patients. Small vessels were congested in most subjects with moderate perivascular extravasation at the level of the medulla, pons and deep cerebellar nuclei in 6 cases. Variable degrees of astrogliosis were evident in all subjects in all assessed regions, but were more pronounced at the level of the brainstem, as testified by GFAP staining (Supplementary Figure 3; semi-quantitative evaluation of astrogliosis across brain regions is available in Supplementary Table 2). Alzheimer Disease (AD) neuropathological changes, evaluated according to NIA-AA criteria, as well as Cerebral Amyloid Angiopathy (CAA) were detected in 5 subjects. In one case, Parkinson’s Disease neuropathological alterations (i.e. deterioration of the ventrolateral substantia nigra and nigral lewy bodies) were found.

Control subjects presented similar macroscopic and histopathological alterations: mild-to-moderate generalized cerebral atrophy (N=7), mild-to-moderate diffuse cerebral edema (N=11) and chronic territorial ischemic injury (N=7); most subjects who died due to pneumonia or respiratory failure presented variable degrees of diffuse hypoxic / ischaemic damage, with mild to moderate damage of the brainstem being a common finding, similarly to COVID-19 subjects (Figure 1C); individual findings for hypoxic / ischaemic injury are reported in Table 2. Four subjects presented AD neuropathological changes and CAA, with one subject presenting both AD and Lewy Body Dementia mixed pathology. The macroscopic and histopathological findings of both COVID-19 subjects and controls are reported in Table 1 and Table 2.

### CNS platelet-enriched microthrombi in small parenchymal vessels were detected in COVID-19 subjects, but not in controls

Small vessel thromboses were detected in 9 COVID-19 patients at the level of the pons, deep cerebellar nuclei and cerebral cortex, with one patient presenting small vessel thromboses in multiple sites. No CNS or systemic thromboses were detected in controls. In all COVID-19 cases, CD61 immunoperoxidase staining revealed platelet-rich microthrombi in small parenchymal vessels, with no evidence of arachnoid of meningeal vessels being involved, as seen in Figure 1D. Other organs were often affected, such as the lungs, liver, intestine, and hypopharynx and even the carotid body^10, 27–28^, as summarized in Table 1. In 3 out of 9 cases, microthromboses were identified only within the CNS, while in the remaining 6 subjects, pulmonary thromboses were also detected. Interestingly, 3 out of 9 subjects with CNS microthrombi were on antithrombotic medication, 4 were not actively treated prior to death, and in 2 cases clinical information regarding antithrombotic medication was incomplete. In line with previous findings in literature, CNS microthromboses appear to be peculiar to the COVID-19 cohort, with no control subject presenting either fibrin- or platelet-enriched microthrombi in the CNS or other organs regardless of the cause of death.

### Microglial cells with an activated phenotype and frequent microglial nodules were found in COVID-19 subjects, but not in controls

In 23 COVID-19 subjects parenchymal microglia displayed an activated phenotype with characteristic thorny ramifications or amoeboid morphology (Figure 2A-B). Interestingly, homeostatic microglial marker TMEM119 was consistently expressed in our cohort (Figure 2A-D), even though it is known to be downregulated upon microglial activation in various neuropathological conditions^36^. A similar pattern of immunoreactivity is also seen in Matschke et al.^8^ and Schwabenland et al.^18^. Considering the relatively short hospitalization time prior to death of our COVID-19 cohort (14.5 days), and the similar immunoreactivity pattern compared to other available studies, it could be inferred that TMEM119 downregulation does not occur early in COVID-19.

While in both COVID-19 subjects and controls microglial marker TMEM119 and lysosomal-activity marker CD68 were found within the same cell (2A-D), COVID-19 subjects displayed a more widespread CD68+ immunoreactivity (2A-B), with statistically significant differences in CD68 immunoreactive area (A%) at the level of the medulla and midbrain, but not the pons, between the two groups (Figure 2E, Welch ANOVA W= 42.68; medulla p<0.0001; pons p=0.733; midbrain p<0.0001). Ki-67 immunoperoxidase staining, as well as Ki-67 / CD68 double label immunofluorescent staining did not reveal significant Ki-67 immunoreactivity ascribable to microglial cells, suggesting local microglial activation and migration without active proliferation in the considered cases. Microglial nodules associated with perineuronal HLA-DR+ / TMEM119+ / CD68+ cells were suggestive of neuronophagia in 18 COVID-19 subjects (Figure 3A-B, 4G, 6F) and were identified at the level of the substantia nigra (N=14), dorsal motor nucleus of the vagus (N=12), medullary reticular formation (N=9), area postrema (N=6) and basal ganglia (N=5); no microglial nodules were found in control cases, regardless of cause of death. Moreover, moderate to severe infiltration of CD68+/TMEM119-cells was found in 23 subjects (2A); given their prominent perivascular localization, these were likely monocyte-derived macrophages.

### In COVID-19 subjects, a topographically defined pattern of microgliosis was found in the medulla oblongata and midbrain

At the level of the medulla oblongata, Welch one-way ANOVA of individual counting fields (FOVs) (Figure 4C, F) revealed statistically significant differences (p<0.001) in TMEM119+/CD68+ activated microglial cells between the medullary tegmentum (T, FOV-13; 216,84±52,26 microglia / mm^2^) and the ventral medulla (pes, P, FOV4-6; 156,09±35,16 microglia / mm^2^) (Figure 4E-F); no differences were found between individual counting fields of the same anatomical compartment. Furthermore, no significant differences were found when comparing microglial density between IOT and non-IOT COVID-19 patients, as well as AD and non-AD patients with SARS-CoV-2 infection (Figure 4A). When comparing microglial density between COVID-19 patients and the control cohort, statistically significant differences were found when considering overall medullary microgliosis, as well as single anatomical compartments (Figure 4C).

At the level of the pons, Welch one-way ANOVA of individual counting fields revealed statistically significant differences only between the most dorsally located counting field comprising the locus coeruleus (FOV1), and other counting fields (FOV2-6) (Figure 5D, C). However, as for the medulla, no differences were found between IOT and non-IOT patients, as well as for AD versus non-AD patients (Figure 5A). Differences in overall microgliosis, as well as differences between anatomical compartments, were not significant between COVID-19 subjects and controls (Figure 5B). Hence, while there appears to be a higher degree of microgliosis in proximity to the locus coeruleus in COVID-19 when compared to other regions of the pons, no differences were found within COVID-19 subgroups and when compared to controls, indicating pontine microgliosis as a non-specific alteration in our cohort (Figure 5E-F).

At the level of the midbrain, COVID-19 subjects presented marked topographical differences between counting fields comprising the substantia nigra (midbrain tegmentum, FOV1-2 and FOV3, Figure 6C) when compared to counting fields of the midbrain tectum and pes (FOV4-6), as seen in Figure 6C-D. This anatomically segregated pattern of inflammation targeting mainly the substantia nigra, but also part of the pre-acqueductal tegmentum, indicates an increasing dorsal-to-ventral gradient of microgliosis which affects the gray matter of the midbrain, sparing counting fields falling within the cerebral peduncle (FOV5-6). Similar to other brainstem levels, no statistically significant differences in overall microgliosis were found when comparing IOT and non-IOT subjects, as well as AD and non-AD subjects (Figure 6A). When compared to controls, COVID-19 subjects presented significantly higher microglial densities when considering both overall microgliosis, as well as microglial densities within anatomical compartments (Figure 6E), suggesting for a COVID-19-specific microglial response at the level of the midbrain (Figure 6F-G).

**Figure 6.**
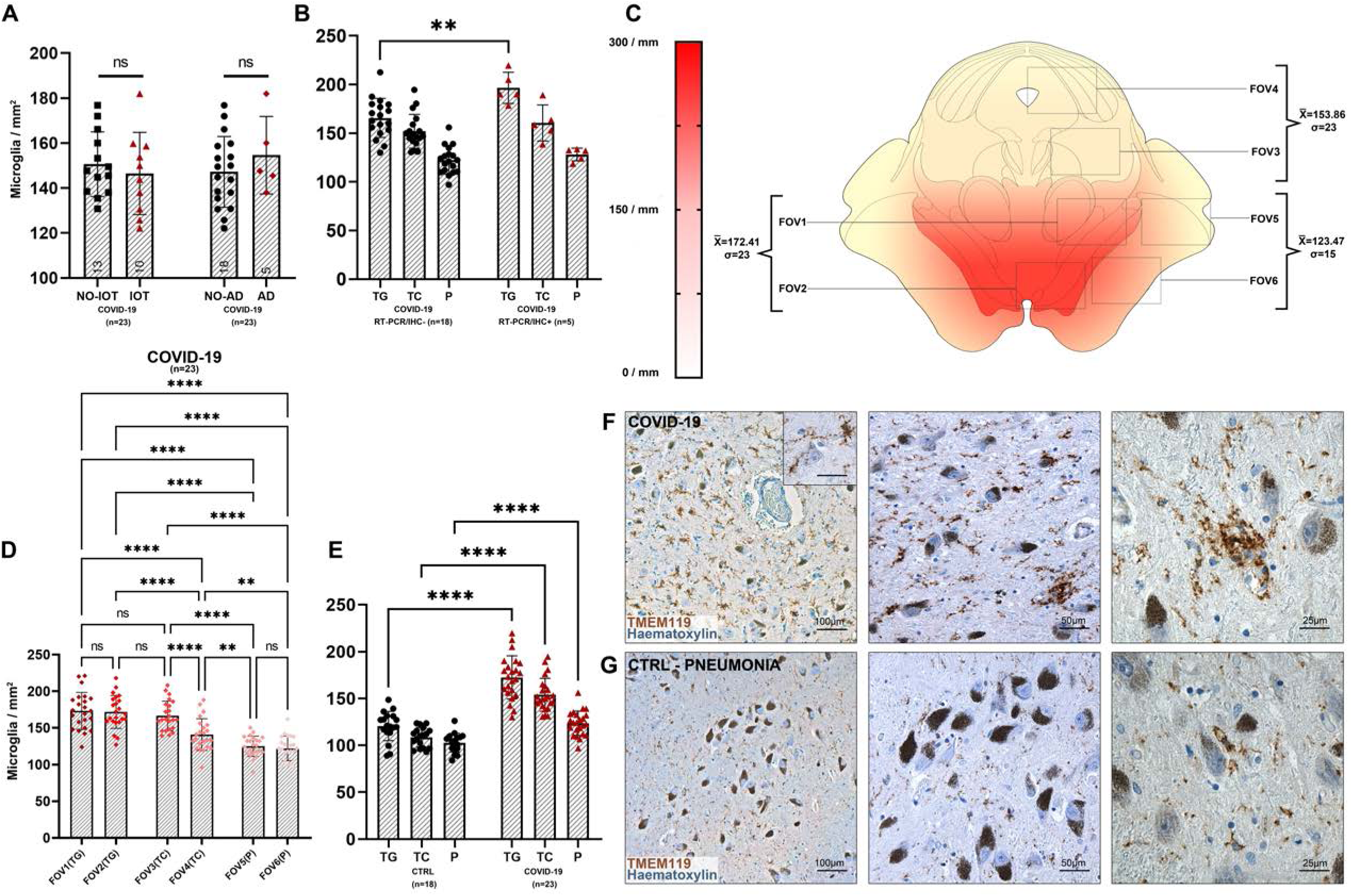
**A)** Welch corrected T-test plot of microglial densities (microglia / mm^2^) in the midbrain of COVID-19 subjects with (n=5, red) and without (n=16, black) Alzheimer’s Disease neuropathological changes (p>0.05), and of COVID-19 subjects treated (n=10, red) and not treated (n=11, black) with intensive oxygen therapy (p>0.05). **B)** Welch one-way ANOVA between anatomical compartments (TG, tegmentum; TC, tectum; P, pes) between COVID-19 subjects with (n=5, red) and without (n=18) viral tropism (RT-PCR/IHC+ versus RT-PCR/IHC-) reveals statistically significant differences at the level of the midbrain tegmentum (p=0.0074), but not other anatomical districts. **C)** Anatomical heatmap of activated microglia within the medulla oblongata in COVID-19. **D)** Welch one-way ANOVA of microglial densities per counting fields (FOV) reveals statistically significant differences between FOVs of the Tegmentum (T; FOV1-2) when compared to FOVs of the Pes (P; FOV5-6), suggesting for a localized pattern of microgliosis comprising the preacqueductal tegmentum and the substantia nigra. **E)** Welch one-way ANOVA between anatomical compartments (TG, tegmentum; TC, tectum; P, pes) between COVID-19 subjects (n=23, red) and controls (n=18, black) reveals statistically significant differences between all anatomical districts of the midbrain (p<0.0001). **F-G)** TMEM119 immunoperoxidase staining of comparable regions of the midbrain in COVID-19 subjects (above) and controls (below). Inset: perineuronal microglia in the substantia nigra. COVID-19 subjects often present distinct microglial nodules.

**Figure 7.**
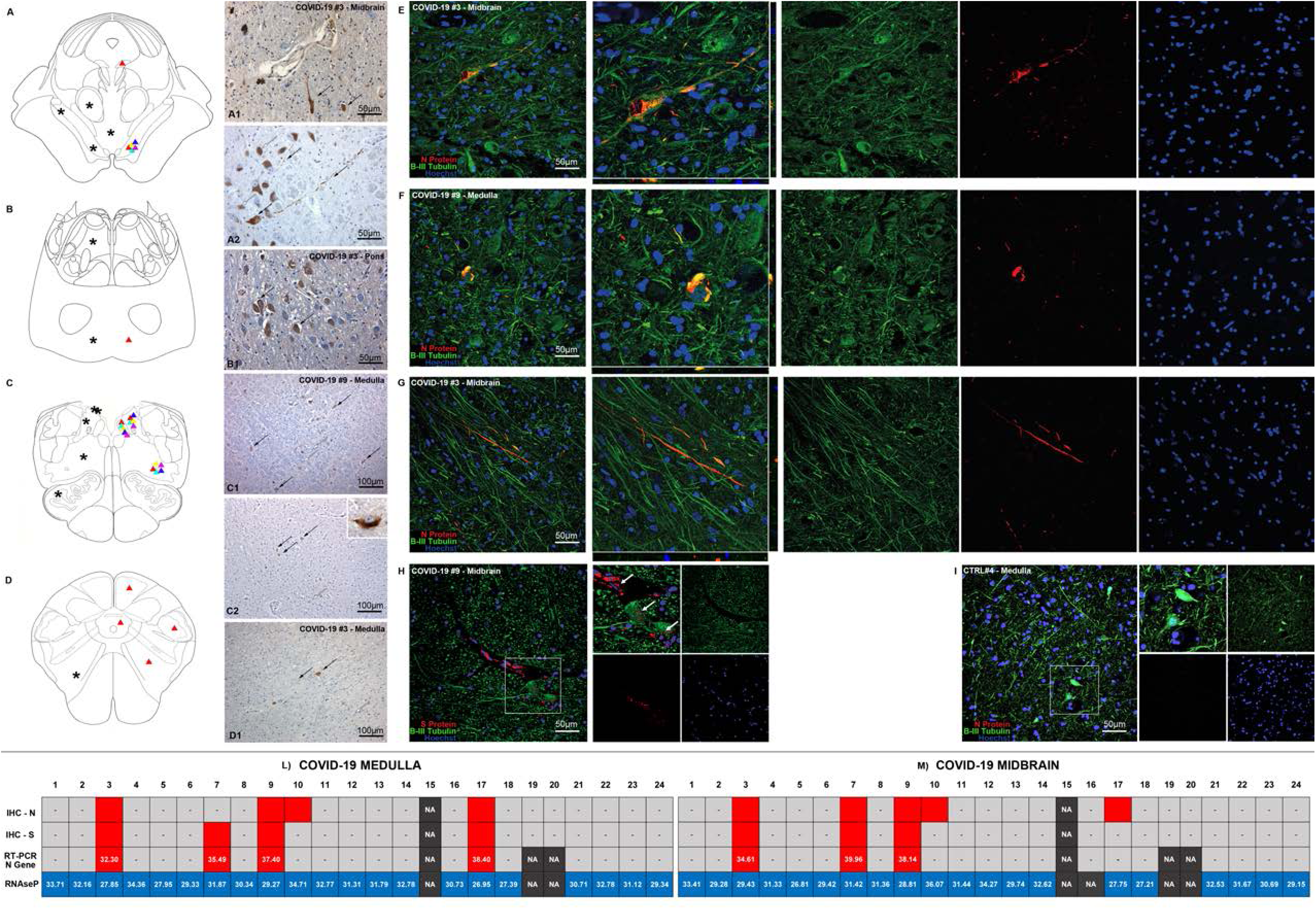
Topographical localization of SARS-CoV-2 Viral Protein Immunoreactivities (Triangles, Right Half) and Microglial Nodules (Asterisks, Left Half) throughout the brainstem. **A)** At the level of the Mesencephalon, Immunoreactivities are found mainly within the boundaries of the substantia nigra, with the exception of Subject #3, which also presented immunoreactive neurons within the Interstitial Nucleus of Cajal; Microglial nodules were confined mainly within the boundaries of the tegmentum, and were not detected neither within the pes nor the tectum. **A1)** SARS-CoV-2 Spike Protein IHC at the level of the Substantia Nigra reveals immunoreactive neurons (mean of 2 immunoreactivities per mm^2^) with well-marked processes (black arrows); negative neurons can also be found nearby (white arrows). **A2)** SARS-CoV-2 Nucleocapsid Protein IHC reveals a similar pattern of immunoreactive neurons and axons throughout the substantia nigra. **B)** At the level of the pons, Subject #3 presented immunoreactive neurons (mean of 5 immunoreactivities per mm^2^) within the basilary nuclei, while microglial nodules were found both within the basis, as well as the dorsal pons in proximity to the facial nucleus. **B1)** SARS-CoV-2 Spike Protein IHC at the level of the pons in Subject #3, displaying immunoreactive neurons (black arrows) within the basilary nuclei of the pons; non-reactive cells can also be appreciated (white arrows) **C)** At the level of the upper medulla oblongata, immunoreactivities were found at the level of the dorsal motor nucleus of the vagus, solitary tract nucleus and nucleus ambiguus; microglial nodules were prominent within the Vagal Trigone and Area Postrema, but were also found within the reticular formation and the inferior olivary complex. **C1-2)** SARS-CoV-2 Spike Protein IHC at the level of the solitary tract nucleus and nucleus ambiguus; immunoreactive neurons can be seen within the anatomical boundaries of these nuclei (black arrows), along with non-reactive cells (white arrows). Inset of a single reactive neuron within the solitary tract nucleus, Spike Protein immunohistochemistry. **D)** At the level of the Lower Medulla Oblongata, Immunoreactivities were found at the level of the spinal trigeminal nucleus and medullary reticular formation. Microglial nodules were found within the medullary reticular formation. **D1)** SARS-CoV-2 Spike Protein IHC at the level of the medullary reticular formation in the lower medulla (black arrows); non-reactive cells are indicated with a white arrow. **E-I)** Double label N and S Protein (red) and Beta-III Tubulin (green) fluorescent immunohistochemistry in COVID-19 subjects (E-H) and controls (I). Distinct immunoreactive neurons and neurites can be appreciated in both the medulla (dorsal motor nucleus of the vagus) and midbrain (substantia nigra) (E-G). Juxtavascular immunoreactive neurons in proximity to an immunoreactive vessel in the midbrain of COVID-19 subject #9 (H, inset). Control subjects present no viral protein immunoreactivity (I). **L-M)** N and S protein IHC, real time RT-PCR Cycle Thresholds for SARS-CoV-2 N Gene and RNAseP quality control in our COVID-19 cohort at the level of the medulla (L) and midbrain (M).

We also found a strong correlation between microglial densities across the different levels of the brainstem, as well as CD68+ A% of the corresponding level, as summarized in Figure 2G. The strong positive correlation between microglial density and CD68 immunoreactive area further underlines the activated phenotype displayed by microglial cells in COVID-19. Interestingly, in the COVID-19 cohort, hospitalization time was positively correlated to microglial density in the medulla (r= 0.44; p=0.044), but not with microglial density in the pons and midbrain (Figure 2F). This appears to indicate an increase of microglial densities in the medulla as infection progresses, while the levels of microgliosis within the rest of the brainstem appear to remain relatively stable throughout time. Considering the numerous instances of microglial nodules and neuronophagia encountered in the medulla of the COVID-19 cohort, this is suggestive of prominent medullary impairment ongoing during COVID-19, regardless of oxygenation status or prior neurodegenerative pathology. Conversely, microglial density in the medulla also correlates with hypoxic / ischaemic damage of the brainstem, evaluated along a four-tiered semi-quantitative scale (r= 0.59; p=0.004), as also seen in Thakur et al.^37^. Hence, while microgliosis is strongly characteristic of COVID-19 subjects and differs from controls, brainstem hypoxia / ischaemia plays a major role in mediating medullary microgliosis, as seen in our cohort and in accordance to previous literature.

### SARS-CoV-2 Viral proteins were detected in neurons of the medulla and midbrain in a subset of COVID-19 subjects, but not in controls

Immunoperoxidase and immunofluorescent staining for SARS-CoV-2 spike protein and nucleocapsid protein was performed on all samples of included subjects, showing only positive results in cases with SARS-CoV-2 infection, but not in controls, indicating specificity. In particular, viral proteins were detected in seven subjects (#3, #7, #9, #10, #11, #17, #18) within CNS parenchyma and in five subjects (#3, #7, #9, #10, #17) with immunoreactive neurons within the anatomically defined boundaries of the solitary tract nucleus, dorsal motor nucleus of the vagus, nucleus ambiguus and substantia nigra (Figure 7A-D). As seen in double immunofluorescence labeling, SARS-CoV-2 Nucleocapsid protein antibody can be detected in β-III Tubulin (a pan-neuronal marker) immunoreactive structures, such as neuronal somata and neurites in the medulla and midbrain (Figure 7E-H), with no labeling in controls (Figure 7I). At the level of the midbrain, Nucleocapsid protein immunofluorescence was also found within tyrosine hydroxylase immunoreactive neurons and neurites of the substantia nigra, indicating the presence of viral antigens within dopaminergic neurons (Figure 8A-E). Some of these subjects (#7, #9, #11, #17, #18) also displayed endothelial cell immunoreactivity in small vessels of the cerebral cortex (subject #11), deep cerebellar nuclei (#17-18) hippocampus (#7) (Supplementary Figure 2) and midbrain (#9) (Figure 7H); small vessel thromboses, perivascular extravasation and hemorrhagic injury were found within affected regions of these cases.

**Figure 8.**
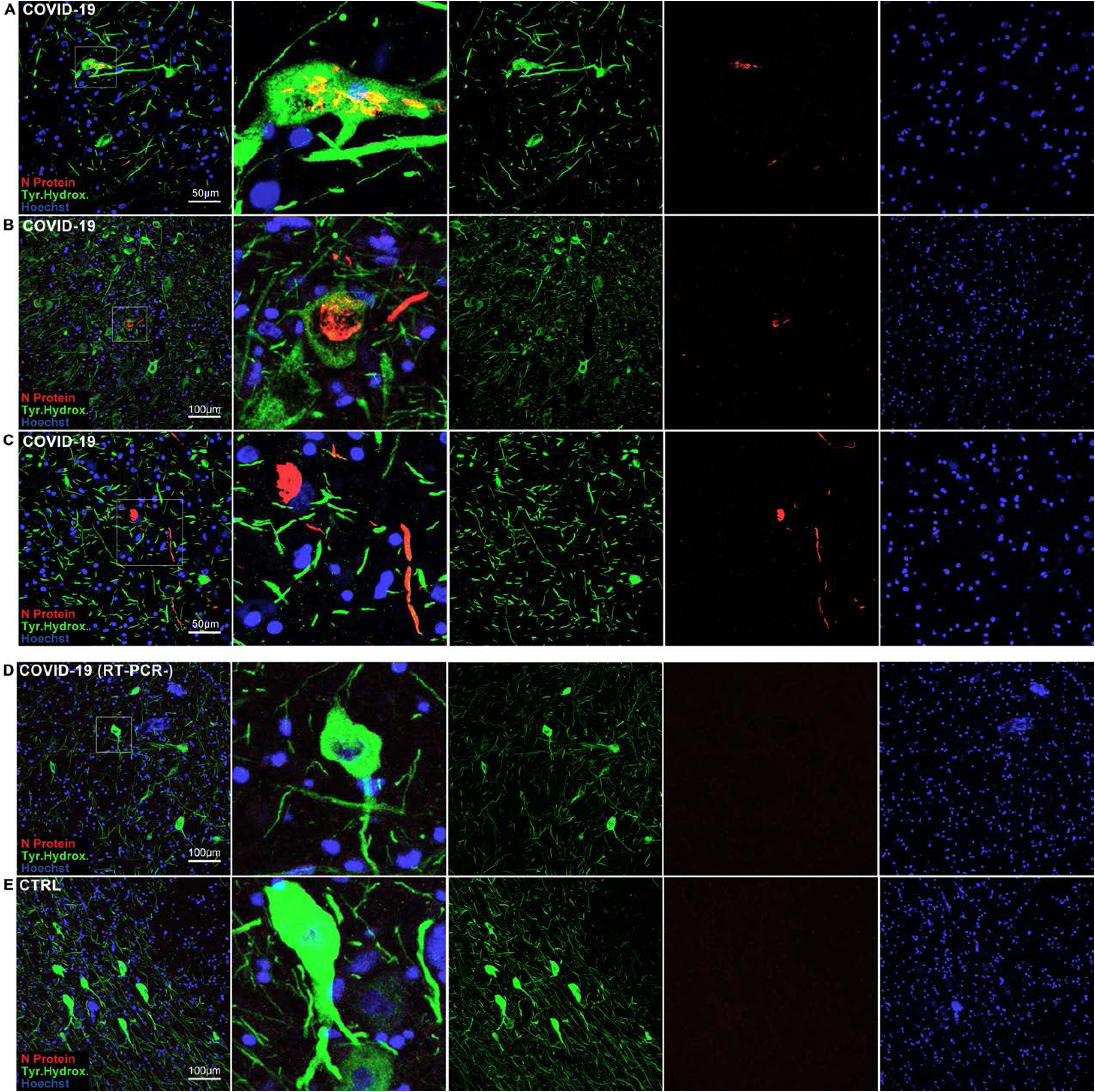
Double label N Protein (red) and Tyrosine Hydroxylase (green) fluorescent immunohistochemistry at the level of the substantia nigra in the midbrain. **A-C)** in COVID-19, Both TH+ and TH-neurons display N protein immunoreactivity. **D-E)** in COVID-19 subjects with negative RT-PCR/IHC (D) as well as non-COVID controls (E) no N protein staining was detected.

In case #7, ischaemic injury of the right rostral hippocampal formation due to posterior cerebral artery (PCA) occlusion was associated with perivascular extravasation, edema, fibrinogen leakage and viral protein immunoreactivity within small vessel endothelium, further confirmed by RT-PCR. Acute hemorrhagic injury in the territory of the right middle cerebral artery (MCA) in #11 (Figure 1B) was associated with endothelitis within perilesional tissue, displaying both viral protein immunoreactive endothelium and positive RT-PCR. Similarly, the deep cerebellar white matter and dentate nuclei in cases #17-18 presented small vessel thromboses and extensive hemorrhagic injury (case #17). Conversely, in some cases with small vessel thromboses within the pons and frontal cortex (e.g. #19-20), viral proteins and RNA was not detectable. There was no correlation between viral protein immunoreactivity / RT-PCR Cycle threshold and hospitalization time (Figure 4E), suggesting no apparent link between the detection of viral antigens and genomic sequences and post-infection interval. However, the actual length of infection, particularly prior to hospitalization, could not always be safely determined, as pre-symptomatic infection could not be excluded, nor evaluated, from the available clinical data. ACE2 receptor protein and TMPRSS2 protein immunoreactivity was compatible with the anatomical distribution of SARS-CoV-2 antigens (Supplementary Figure 1, E-F), as detected with immunoperoxidase staining, but was not consistently replicated in immunofluorescent staining (data not shown). Both proteins were moderately expressed in vascular endothelial cells and brainstem neurons.

### RT-PCR analyses of FFPE tissue sections detected viral RNA in COVID-19 cases with viral protein immunoreactivity

Molecular testing by real-time RT-PCR detected SARS-CoV-2 RNA in 10 out of 24 COVID-19 subjects, 9 of whom had also SARS-CoV-2 S and/or N protein-positive IHC / IF (Figure 7L-M, Supplementary Table 2). In positive tissue samples, threshold cycles (Ct) of real-time RT-PCR for SARS-CoV-2 RNA ranged between 33 and 38, while in all samples the Ct values of the internal control RNAseP ranged between 27 and 34. The cycle threshold values for each analyzed section are reported in Supplementary table 2 and in Figure 7L-M for the medulla and midbrain. SARS-CoV-2 subgenomic RNA was investigated but not detected in our specimens, likely due to RNA degradation within FFPE sections, as indicated by the low Ct values of RNAse P.

### Viral antigens are associated to higher microglial densities within affected anatomical loci, but no differences are found in overall microgliosis, suggesting a specific topographical response

While overall levels of microgliosis within the medulla, pons and midbrain did not differ significantly between COVID-19 subjects with and without detectable viral antigens, Welch one-way ANOVA between anatomical compartments (i.e. tegmentum, tectum and pes) revealed statistically significant differences within the COVID-19 cohort. Indeed, subjects with detectable viral genomic sequences and antigens (RT-PCR+/IHC+) were characterized by higher microglial densities in the medullary (p=0.017) and midbrain tegmentum (p=0.0074) when compared to negative (RT-PCR-/IHC-) COVID-19 subjects, as seen in Figure 4B and 6B. In association with frequent instances of microglial nodules and perineuronal TMEM119+/CD68+ microglial cells suggestive of neuronophagia (Figure 3A-B), this finding suggests a peculiar microglial response towards anatomical loci of the brainstem in which SARS-CoV-2 antigens were detected, even though overall levels of microgliosis within brainstem regions did not appear to differ significantly. Taken together with little-to-no Ki-67 immunoreactivity and no detectable Ki-67+ / CD68+ immunofluorescent signal, migration of microglial cells towards the site of injury appears to be the more likely mechanism occurring in COVID-19 inflammation, rather than microglial proliferation within affected regions.

## Discussion

In the present study, the neuropathological findings of 24 COVID-19 patients were examined and compared with age- and sex-matched controls who died due to pneumonia and / or respiratory insufficiency. Our findings indicate, in line with some of the previous autopsy reports, specific neuropathological alterations in the brains of COVID-19 patients, with particular regard to topographically-defined microgliosis within anatomical loci of the brainstem and viral immunoreactivity in specific CNS compartments, either within the boundaries of brainstem nuclei or in the context of ischaemic and hemorrhagic injuries. Platelet and fibrin microthrombi, in particular, were characteristic findings of the COVID-19 cohort, and often affected multiple organs, such as the lungs, liver, intestine, hypopharynx and even the carotid body^10, 27–28^, as summarized in Table 1. Microthromboses were more frequent within the pons, deep cerebellar nuclei and cerebral cortex. In some cases, hemorrhagic injury and microthromboses were found in regions with viral protein immunoreactivity in vascular endothelial cells.

SARS-CoV-2 viral antigens, on the other hand, were confined to specific loci of the CNS. As seen in Figure 7A-H, SARS-CoV-2 appears to be localized preferentially within neurons of the vagal nuclei of the medulla and the substantia nigra, with the exception of one subject who also presented immunoreactive cells throughout the whole brainstem (#3). While Matschke et al.^8^ reported SARS-CoV-2 invasion of cranial nerves IX-X, we were unable to replicate these findings within our cohort; furthermore, unlike Meinhardt et al.’s findings^9^, viral proteins and RNA were not detectable in any of the sampled olfactory bulbs, tracts and bifurcations, even though moderate edema, moderate-to-severe astrogliosis and moderate microglial activation was encountered in most cases in our study. ACE2 Receptor and TMPRSS-2 protein immunohistochemistry support this topographical localization, with neurons within the dorsal motor nucleus of the vagus, solitary tract nucleus, nucleus Ambiguus and Substantia Nigra being moderately immunoreactive (Supplementary Figure 1, E-F).

While previous studies identified viral protein immunoreactivity in sparse cells throughout the brainstem^8–18^ without specific topography, our findings appear to be in line with available animal studies on other coronaviruses, i.e. SARS-CoV and MERS-CoV, which are known to be able to infect the brainstem, and particularly the dorsal motor nucleus of the vagus, solitary tract nucleus and nucleus ambiguus, so that an analog pattern of neuroinvasion for SARS-CoV-2 has been suggested^17, 29–31^. The peculiar and unexpected finding in our cohort was the detection of viral antigens and genomic sequences within the substantia nigra, not matching any known models of coronavirus neurotropism. Interestingly, SARS-CoV-2 S and N protein were detected in both tyrosine hydroxylase positive and negative neurons (Figure 8). Immunoreactive neurons of the substantia nigra were frequently found in proximity to blood vessels, which were at times immunoreactive to viral proteins as well (Figure 7H, Supplementary Figure 2). Hence, aside from olfactory-transmucosal transmission identified by Meinhardt et al.^9^, and vagus/glossopharyngeal-mediated invasion identified by Matshke et al.^8^, SARS-CoV-2 may gain access to other districts of the CNS either through a yet-unknown neuronal route or, as suggested by our findings in the midbrain and as seen in Schwabeland et al.’s study^18^, by crossing the blood-brain-barrier and infecting structures of the peri- and juxtavascular compartment. We believe these findings encourage further research on the possibility that these events may be the trigger of a neurodegenerative process such as Parkinson disease in susceptible individuals. Future studies on COVID-19 survivors and Long COVID patients are therefore warranted^39^.

However, despite the detection of viral proteins and genomic sequences in restricted regions of the brainstem, we found no evident neuropathological alterations in SARS-CoV-2 infected cells, such as necrotic changes and other cytological alterations, that could hint towards possible direct consequences of viral invasion in human neurons. COVID-19 is characterized by different evolutionary phases and heterogeneous individual responses, and the short interval between infection and death in our cohort (mean hospitalization time = 14 days), as well as the fact that included patients died during the acute phase of the disease, may not be sufficient to determine detectable neuropathological alterations in affected cells as a direct consequence of viral invasion, which may require more time to develop^3, 31^. Lastly, the detection of viral proteins in a subset of patients (5 out of 24), as seen in this and previous studies may be related to the particularly severe disease, and concurring comorbidities, of the patients subject to neuropathological examination.

Hence, while the consequences of SARS-CoV-2 neurotropism in the medulla have been widely discussed in literature, and are supported by the detection of viral proteins and genomic sequences in our study, the absence of direct neuronal damage and the impossibility of performing functional assays on post-mortem samples should be taken into consideration when discussing the clinical implications of COVID-19 neuropathology. Future studies on “long COVID” patients^3^ may be able to shed a light on the long-term consequences of COVID-19, particularly concerning the detection of SARS-CoV-2 within the CNS after the acute phase of the disease, and whether or not this leads to specific neuropathological alterations as a consequence of viral invasion.

Concerning microglial activation and density, our findings appear to be in line with Schwabenland et al.^18^, who identified microglial nodules and parenchymal reactive microglia as hallmark for COVID-19, in contrast to both controls and ExtraCorporeal Membrane Oxygenation (ECMO) patients. In our cohort, patients with pneumonia and / or respiratory failure served as control group and, although also characterized by microglial activation, displayed lower microglial counts in the medulla and midbrain, but not in the pons, when compared to COVID-19 subjects. We also found no significant effect of oxygen therapy on microglial density within the COVID-19 group. Conversely, Deigendesh et al.^33^ found significant differences in HLA-DR+ activated microglia when comparing COVID-19 subjects to non-septic controls, but no differences were found with patients who had died under septic conditions; according to the authors^33^ this may represent a histopathological correlate of critical illness-related encephalopathy, rather than a COVID-19-specific finding. Aside from the distinct populations serving as control subjects, significant methodological differences between these studies must be taken into consideration. Our approach to microglial quantification was more similar to Schwabenland et al.^18^, as digitally-assisted manual counting of TMEM119+ cells, a homeostatic microglia-specific marker, was performed to estimate microgliosis; however, we also expanded on these findings by estimating microgliosis in a topographically dependent manner by employing set counting fields within anatomically defined regions; conversely Deigendesh et al.^33^ quantified HLA-DR immunoreactive area, a marker expressed on both microglia and on infiltrating lympho-monocytic cells, as a fraction of the counting field (A%), and not as individual particles. Hence, while our estimation selectively reflects the activity of microglial cells in COVID-19 and pneumonia controls within anatomically defined regions of the brainstem, Deigendesh’s study offers a broader representation of overall brainstem inflammation under different conditions, including patients deceased under septic conditions, explaining differences between our studies.

Interestingly, while no evidence of direct neuronal damage was found in SARS-CoV-2 infected cells, microglial densities within affected anatomical loci differed between subjects with and without detectable viral antigens and genomic sequences (RT-PCR/IHC+ versus RT-PCR/IHC-in Figure 4B and 6B), suggesting a link between the detection of SARS-CoV-2 antigens and microglial response. Conversely, overall microglial density (i.e. without topographical delineation) did not differ between the two groups, and a strong correlation between microgliosis and hypoxic / ischaemic damage at the level of the brainstem was found. Hence, while we found a suggestive link between microgliosis and the detection of SARS-CoV-2 antigens in our cohort, other factors such as hypoxia / ischaemia and systemic inflammation / cytokine storm ongoing during COVID-19, as previously reported by Thakur et al^37^, are likely to play a more prominent role in determining brainstem microgliosis, in accordance to previous studies^33^.

## Conclusions

The present study contributes to define the spectrum of neuropathological alterations in COVID-19, as well as the neuroinvasive potential of SARS-CoV-2 within the CNS. Unlike previous findings, we have documented a subset of COVID-19 cases in which viral proteins and genomic sequences were detectable within anatomically defined regions of the CNS. Similarly, microglial activation in the brainstem appears to differ between COVID-19 and pneumonia / respiratory failure controls, with the former also presenting a pattern of increased microglial density in specific compartments of the medulla and midbrain. However, despite this evidence supporting the neuroinvasive potential of SARS-CoV-2, neuropathological alterations encountered in our cohort cannot be ascribed to viral antigens detected in the brainstem. In line with other studies in literature, hypoxic / ischaemic damage and systemic inflammation likely represent major contributors in determining neuropathological alterations in COVID-19, with little-to-no evidence indicating direct viral damage of the central nervous system in humans. Moreover, further investigation is required to determine whether or not SARS-CoV-2 neurotropism represents a major component of COVID-19 in the general population, as subjects included in neuropathological studies often present a much more severe course of the disease and major medical comorbidities. Nevertheless, the findings of our study suggest the possibility that, although not frequently, SARS-CoV-2 may gain access to specific regions of the central nervous system, especially the vagal nuclei of the medulla and the substantia nigra in the midbrain. As direct neuropathological alterations determined by SARS-CoV-2 neurotropism may not be detectable in subjects deceased during the acute phase of the disease, future studies are required to determine whether or not SARS-CoV-2 neurotropism is present in chronic COVID-19 patients, or in COVID-19 survivors suffering from the long-term effects of infection, and if eventual neuropathological alterations in these subjects can be ascribed to viral tropism, rather than immune-mediated mechanisms.

### Limitations of the study

This study is based on post-mortem tissue samples obtained during the first wave of the COVID-19 pandemic in Italy. While the neuropathological alterations encountered in our work contribute to define the pathological mechanisms of COVID-19 and SARS-CoV-2 infection in the CNS, the lack of exhaustive post-infection neurological evaluation of included patients does not allow for unequivocal clinico-pathological correlations. It must also be considered that most patients included in the study died during the peak of the sanitary emergency in Italy, one of the first countries to face the COVID-19 pandemic in Europe, and neurological evaluation was not always possible. Hence, it remains to be determined whether the neuropathological alterations observed in this study are also linked to neurological symptoms, and whether they are also present in COVID-19 survivors.

Unlike previous studies in literature, we have included 18 controls who died due to pneumonia, respiratory insufficiency or multiorgan failure, rather than healthy controls. Retrospective selection of control subjects, however, could lead to unwanted selection bias. Furthermore, from the available clinical data of our controls, we have found no instances of intensive oxygen therapy or mechanical ventilation, but incompleteness of available clinical records cannot entirely be excluded. For this purpose, we have also performed comparisons within the COVID-19 group, identifying no statistically significant differences between subjects with and without neurodegenerative conditions, and no influence of oxygen therapy on brainstem microgliosis. The involvement of other brain regions, such the cerebral and cerebellar cortex and the basal ganglia, cannot be excluded but is beyond the scopes of this study. Moreover, as all patients died during the first wave of the COVID-19 pandemic, our findings may not reflect the possible neuropathological alterations encountered in patients affected by SARS-CoV-2 variants.

Limitations to viral antigen / RNA detection in our study must also be considered. Real-time RT-PCR cannot exclude detection of viral RNA in blood vessels within samples. While particular care was taken to avoid contamination by employing sterile instruments and disposable microtome blades when sampling FFPE sections for RT-PCR analyses, the main strength of our study was the complementary use of immunoperoxidase and immunofluorescent staining with different antibodies to detect viral antigen as an indicator of viral tropism. This is further strengthened by the strong concordance between these assays in our cohort, quantified by a statistically significant positive correlation between RT-PCR cycle threshold and IHC positivity (r=0.87, p<0.0001).

In conclusion, further investigation is required to determine the direct effects of viral invasion within the CNS, with particular regard to cases of long-lasting infection and in COVID-19 survivors.

## Acknowledgements

We are grateful to Prof. James E. Goldman for his feedback and suggestions concerning our study.

## Data Sharing

data is available from the corresponding author upon request; raw data underlying graphs and statistical analyses has been provided to the editor, reviewers, and is available upon reasonable request.

## Ethical approval

All procedures were carried out in accordance to the Declaration of Helsinki. Samples were anonymous to the investigators and used in accordance with the lives of the Committee of the Ministers of EU member states on the use of samples of n origin for research.

