## Supplementary for "COVID-19 Neuropathology: evidence for SARS-CoV-2 invasion of Human Brainstem Nuclei"

**SUPPLEMENTARY TABLE 1.** Anatomical boundaries and main structures contained in each examined Field of View, divided per level of sectioning.

| LEVEL OF SECTIONING | **FOV1** | **FOV2** | **FOV3** | **FOV4** | **FOV5** | **FOV6** |
|---|---|---|---|---|---|---|
| MEDULLA | Tegmentum: Dorsal Motor Nucleus of the Vagus, Solitary Tract Nucleus, Area Postrema | Tegmentum: Hypoglossal Nucleus, Intercalate Nucleus, Medial Longitudinal Lemniscus, Dorsal Medullary Reticular Formation | Tegmentum: Nucleus Ambiguus, Ventral Medullary Reticular Formation. | Basis: Medial Lemniscus, Medial Accessory Olivary Nucleus, Inferior Olivary Complex (Medial Part) | Basis: Inferior Olivary Complex (Lateral Part). | Basis: Pyramidal Tract. |
| PONS | Tegmentum: Dorsal Tegmentum, Locus Coeruleus, Medial Longitudinal Fasciculus. | Tegmentum: Ventrolateral Tegmentum, Spinal Trigeminal Tract. | Tegmentum: Ventromedial Tegmentum, Medial lemniscus, Central Tegmental Tract. | Basis: Dorsomedial basis pontis, basilary nuclei of the pons, cortico-spinal tract. | Basis: Lateral basis pontis, basilary nuclei of the pons, cortico-spinal tract. | Basis: Ventromedial basis pontis, basilary nuclei of the pons, cortico-spinal tract. |
| MIDBRAIN | Tegmentum: Lateral part of the Substantia Nigra, Red Nucleus | Tegmentum: Medial part of the Substantia Nigra. | Tectum: Quadrigeminal Plate, Periacqueductal Gray. | Tegmentum / Tectum interface, Mesencephalic reticular formation. | Basis: lateral part of the cerebral peduncle. | Basis: medial part of the cerebral peduncle. |

**SUPPLEMENTARY TABLE 2.** Histopathological and Immunohistochemical findings in the COVID-19 Subjects. N and S indicate the viral antigen detected in the tissue. N, Nucleocapsid Protein; S, Spike Protein; RT-CT, Real Time Cycle.

| | |
|---|---|
| | Mild |
| | Moderate |
| | Severe |
| | Absent / Negative |
| | Present / Positive |

| CASE | SAMPLE / LEVEL | ASTROGLIOSIS (GFAP) | MICROGLIOSIS (TMEM119-CD68-HLA-DR) | THROMBOSES / SMALL VESSEL THROMBOSES (H&E – CD61) | VIRAL PROTEINS (IHC) | VIRAL RNA (RT-PCR) |
|---|---|---|---|---|---|---|
| **#1** | Medulla Oblongata | | | | | |
| | Pons | | | | | |
| | Midbrain | | | | | |
| | Cerebellar Cortex | | | | | |
| | Basal Ganglia | | | | | |
| | Frontal Cortex | | | | | |
| | Olfactory bulbs and tracts | | | | | |
| | Cranial Nerves (III-XI) | | | | | |

| | | | | | |
|---|---|---|---|---|---|
| | Leptomeninges | | | | |
| | Choroid Plexuses | | | | |
| **#2** | Medulla Oblongata | | | | |
| | Pons | | | | |
| | Midbrain | | | | |
| | Cerebellar Cortex | | | | |
| | Basal Ganglia | | | | |
| | Frontal Cortex | | | | |
| | Olfactory bulbs and tracts | | | | |
| | Cranial Nerves (III-XI) | | | | |
| | Leptomeninges | | | | **S** |
| | Choroid Plexuses | | | | |
| **#3** | Medulla Oblongata | | | | **S, N** |
| | Pons | | | | **S, N** |
| | Midbrain | | | | **S, N** |
| | Cerebellar cortex | | | | |
| | Basal Ganglia | | | | **S** |

| | | | | | |
|---|---|---|---|---|---|
| | Frontal Cortex | | | | |
| | Olfactory bulbs and tracts | | | | |
| | Cranial Nerves (III-XI) | | | | |
| | Leptomeninges | | | | S |
| | Choroid Plexuses | | | | |
| **#4** | Medulla Oblongata | | | | |
| | Pons | | | | |
| | Midbrain | | | | |
| | Cerebellar Cortex | | | | |
| | Basal Ganglia | | | | |
| | Frontal Cortex | | | | |
| | Olfactory bulbs and tracts | | | | |
| | Leptomeninges | | | | |
| | Choroid Plexuses | | | | |
| **#5** | Medulla Oblongata | | | | |
| | Pons | | | | |

| | |
|---|---|
| | Midbrain |
| | Cerebellar Cortex |
| | Deep cerebellar nuclei |
| | Basal Ganglia |
| | Hippocampus |
| | Frontal Cortex |
| | Olfactory bulbs and tracts |
| | Leptomeninges |
| | Choroid Plexuses |
| **#6** | Medulla Oblongata |
| | Pons |
| | Midbrain |
| | Cerebellar Cortex |
| | Deep cerebellar nuclei |
| | Basal Ganglia |
| | Hippocampus |

| | | | | | | |
|---|---|---|---|---|---|---|
| | Frontal Cortex | | | | | |
| | Leptomeninges | | | | | |
| | Choroid Plexuses | | | | | |
| **#7** | Medulla Oblongata | | | | **S** | RT-CT=35,46 |
| | Pons | | | | | |
| | Midbrain | | | | **S** | RT-CT=39,96 |
| | Cerebellar Cortex | | | | | |
| | Deep cerebellar nuclei | | | | | |
| | Basal Ganglia | | | | | |
| | Hippocampus | | | | **S, N** | RT-CT=38,04 |
| | Frontal Cortex | | | | | |
| | Leptomeninges | | | | | |
| | Choroid Plexuses | | | | | |
| **#8** | Medulla Oblongata | | | | | |
| | Pons | | | | | |
| | Midbrain | | | | | |
| | Cerebellar Cortex | | | | | |

| | | | | | | |
|---|---|---|---|---|---|---|
| | Deep cerebellar nuclei | | | | | |
| | Basal Ganglia | | | | | |
| | Hippocampus | | | | | |
| | Frontal Cortex | | | | | |
| | Leptomeninges | | | | | |
| | Choroid Plexuses | | | | | |
| **#9** | Medulla Oblongata | | | | **S, N** | RT-CT=37,4 |
| | Pons | | | | | |
| | Midbrain | | | | **N** | RT-CT=38,14 |
| | Cerebellar Cortex | | | | | |
| | Deep cerebellar nuclei | | | | | |
| | Basal Ganglia | | | | | |
| | Hippocampus | | | | | |
| | Frontal Cortex | | | | | |
| | Leptomeninges | | | | **S** | RT-CT=37,4 |
| | Choroid Plexuses | | | | | |
| **#10** | Medulla Oblongata | | | | **N** | ND |

| | | | | | | |
|---|---|---|---|---|---|---|
| | Pons | | | | | |
| | Midbrain | | | | **N** | ND |
| | Cerebellar Cortex | | | | | |
| | Deep cerebellar nuclei | | | | | |
| | Basal Ganglia | | | | | |
| | Frontal Cortex and subcortex | | | | **S, N** | RT-CT=37,61 |
| | Leptomeninges | | | | | |
| | Choroid Plexuses | | | | | |
| **#11** | Medulla Oblongata | | | | | |
| | Pons | | | | | |
| | Midbrain | | | | | |
| | Cerebellar Cortex | | | | | |
| | Deep cerebellar nuclei | | | | | |
| | Basal Ganglia | | | | | |
| | Frontal Cortex | | | | | |
| | Parietal Cortex | | | | **S** | RT-CT=38,00 |

| | |
|---|---|
| | Leptomeninges |
| | Choroid Plexuses |
| **#12** | Medulla Oblongata |
| | Pons |
| | Midbrain |
| | Cerebellar Cortex |
| | Deep cerebellar nuclei |
| | Basal Ganglia |
| | Frontal Cortex |
| | Leptomeninges |
| | Choroid Plexuses |
| **#13** | Medulla Oblongata |
| | Pons |
| | Midbrain |
| | Cerebellar Cortex |
| | Deep cerebellar nuclei |
| | Basal Ganglia |

| | |
|---|---|
| | Frontal Cortex |
| | Leptomeninges |
| | Choroid Plexuses |
| **#14** | Medulla Oblongata |
| | Pons |
| | Midbrain |
| | Cerebellar Cortex |
| | Deep cerebellar nuclei |
| | Basal Ganglia |
| | Frontal Cortex |
| | Leptomeninges |
| | Choroid Plexuses |
| **#15* Poor tissue quality** | Medulla Oblongata |
| | Pons |
| | Midbrain |
| | Deep cerebellar nuclei |
| | Basal Ganglia |

| | | | | | | |
|---|---|---|---|---|---|---|
| | Frontal Cortex | | | | | |
| | Leptomeninges | | | | | |
| | Choroid Plexuses | | | | | |
| **#16** | Medulla Oblongata | | | | | |
| | Pons | | | | | |
| | Midbrain | | | | | |
| | Cerebellar Cortex | | | | | |
| | Deep cerebellar nuclei | | | | | |
| | Basal Ganglia | | | | | RT-CT=37,47 |
| | Frontal Cortex | | | | | |
| | Leptomeninges | | | | | |
| | Choroid Plexuses | | | | | |
| **#17** | Medulla Oblongata | | | | **S** | RT-CT=38,34 |
| | Pons | | | | | |
| | Midbrain | | | | **N** | ND |
| | Cerebellar Cortex | | | | | |
| | Deep cerebellar nuclei | | | | **S, N** | ND |

| | | | | | | |
|---|---|---|---|---|---|---|
| | Basal Ganglia | | | | | |
| | Frontal Cortex | | | | | |
| | Leptomeninges | | | | | |
| | Choroid Plexuses | | | | | |
| **#18** | Medulla Oblongata | | | | | |
| | Pons | | | | | |
| | Midbrain | | | | | |
| | Cerebellar Cortex | | | | | |
| | Deep cerebellar nuclei | | | | **S** | RT-CT=37,33 |
| | Basal Ganglia | | | | | |
| | Frontal Cortex | | | | | |
| | Leptomeninges | | | | | |
| | Choroid Plexuses | | | | | |
| **#19** | Pons | | | | | |
| | Midbrain | | | | | |
| | Cerebellar Cortex | | | | | |
| | Deep cerebellar nuclei | | | | | |

| | |
|---|---|
| | Basal Ganglia |
| | Occipital Cortex |
| | Frontal Cortex |
| | Parietal Cortex |
| | Temporal Cortex |
| **#20** | Pons |
| | Midbrain |
| | Cerebellar Cortex |
| | Deep cerebellar nuclei |
| | Basal Ganglia |
| | Occipital Cortex |
| | Frontal Cortex |
| | Parietal Cortex |
| | Temporal Cortex |
| **#21** | Medulla Oblongata |
| | Pons |
| | Midbrain |

| | |
|---|---|
| | Cerebellar Cortex |
| | Deep cerebellar nuclei |
| | Basal Ganglia |
| | Hippocampus |
| | Frontal Cortex |
| | Parietal Cortex |
| | Temporal Cortex |
| | Leptomeninges |
| | Choroid Plexuses |
| **#22** | Medulla Oblongata |
| | Pons |
| | Midbrain |
| | Cerebellar Cortex |
| | Deep cerebellar nuclei |
| | Basal Ganglia |
| | Hippocampus |
| | Frontal Cortex |

| | |
|---|---|
| | Parietal Cortex |
| | Temporal Cortex |
| | Leptomeninges |
| | Choroid Plexuses |
| **#23** | Medulla Oblongata |
| | Pons |
| | Midbrain |
| | Cerebellar Cortex |
| | Deep cerebellar nuclei |
| | Basal Ganglia |
| | Hippocampus |
| | Frontal Cortex |
| | Parietal Cortex |
| | Temporal Cortex |
| | Leptomeninges |
| | Choroid Plexuses |
| **#24** | Medulla Oblongata |

| | |
|---|---|
| | Pons |
| | Midbrain |
| | Cerebellar Cortex |
| | Deep cerebellar nuclei |
| | Basal Ganglia |
| | Hippocampus |
| | Frontal Cortex |
| | Parietal Cortex |
| | Temporal Cortex |
| | Leptomeninges |
| | Choroid Plexuses |

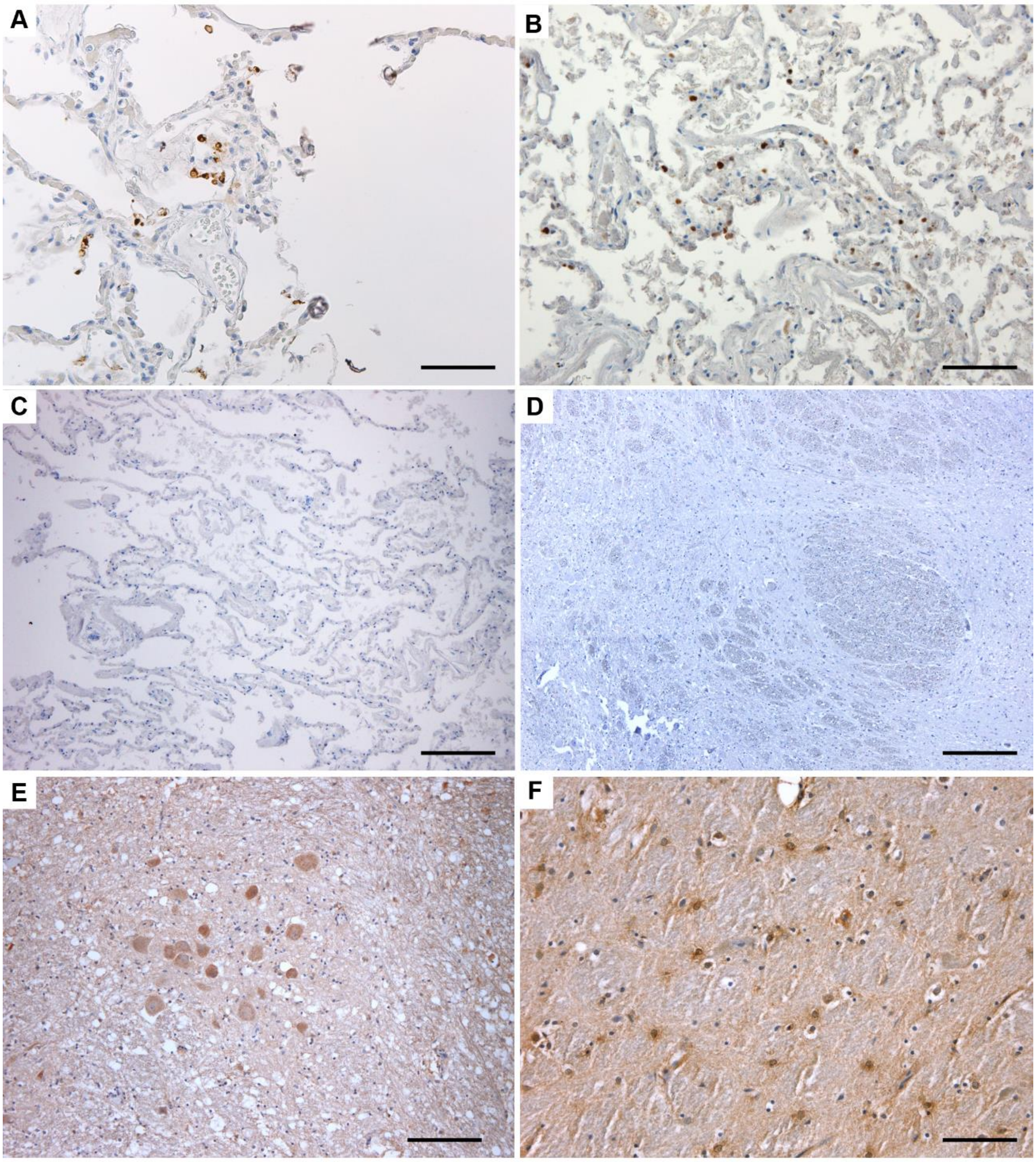

**Supplementary Figure 1. A)** SARS-CoV-2 Spike Protein immunohistochemistry on RT-PCR+ lung tissue. **B)** S SARS-CoV-2 Nucleocapsid Protein immunohistochemistry on RT-PCR+ lung tissue. **C)** SARS-CoV-2 Spike Protein immunohistochemistry on lung tissue from control subjects, showing no immunoreactive elements. **D)** SARS-CoV-2 Spike Protein immunohistochemistry on brainstem tissue (solitary tract nucleus) from control subjects, showing no immunoreactive cells. **E)** ACE-2 Receptor immunohistochemistry at the level of the nucleus ambiguus in the medulla oblongata of a COVID-19 subject, showing diffuse immunoreactivity in the soma and surrounding neuropilum. **F)** TMPRSS-2 protein immunoreactivity in the mesencephalic tegmentum showing moderate reactivity of non-neuronal cells. *Scale Bars: 200µm (C,D); 100µm (A,B,E,F).*

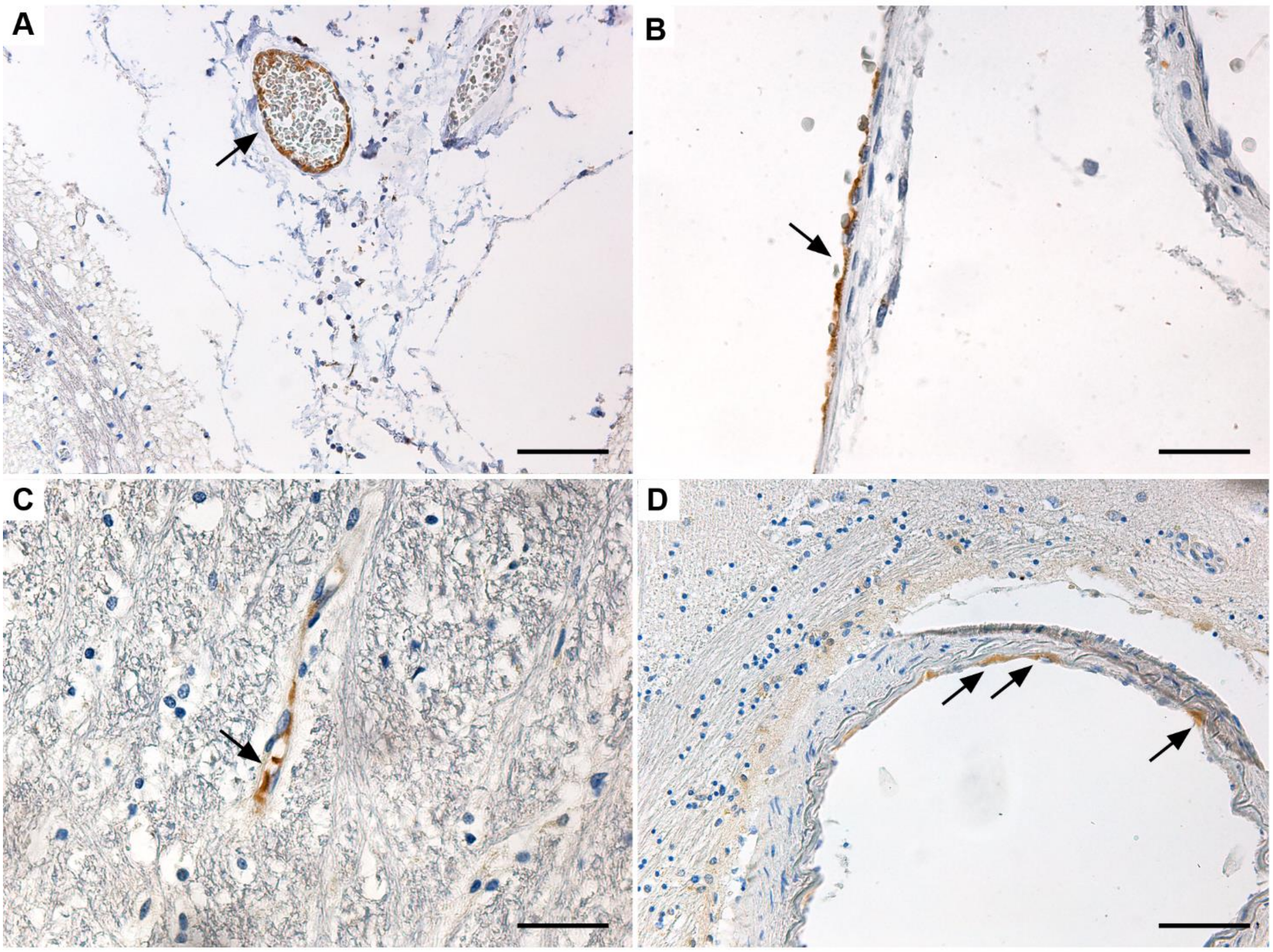

**Supplementary Figure 2. A)** SARS-CoV-2 Spike Protein immunohistochemistry revealing endothelial cell reactivity in a leptomeningeal vessel of the medulla oblongata. **B)** SARS-CoV-2 Nucleocapsid Protein immunohistochemistry in a leptomeningeal vessel in the cerebellum. **C-D)** SARS-CoV-2 Spike Protein immunohistochemistry revealing endothelial cell immunoreactivity in the CNS parenchyma. *Scale Bars: 50µm (A,D); 25µm (B,C).*

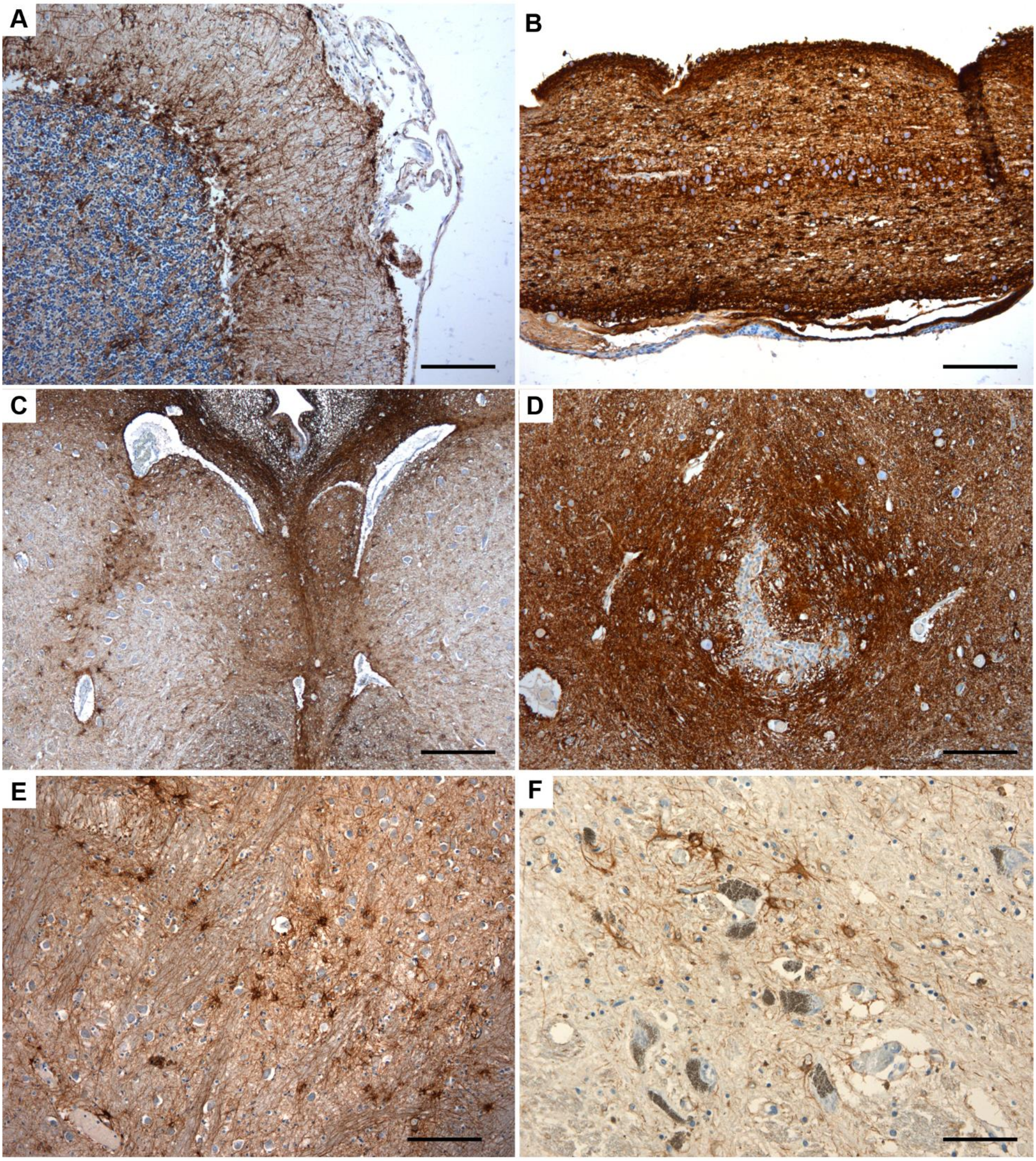

**Supplementary Figure 3.** GFAP Immunohistochemistry. **A)** Reactive Bergmann Glia at the level of the cerebellar cortex in Subject #2. **B)** Marked astrogliosis with numerous corpora amylacea could be appreciated in this sagittal section of the olfactory tract. **C)** Reactive astrocytosis was marked within the subependymal regions of the medullary tegmentum, also displaying numerous reactive astrocytes within the hypoglossal nucleus. **D)** Reactive Astrocytosis surrounding the central canal of the lower medulla. **E)** Reactive astrocytes within the basilary pons in Subject #3. **F)** Reactive astrocytes at the level of the substantia nigra in the midbrain. *Scale Bars: 200µm (C); 100µm (A,B,D,E); 50µm (F).*

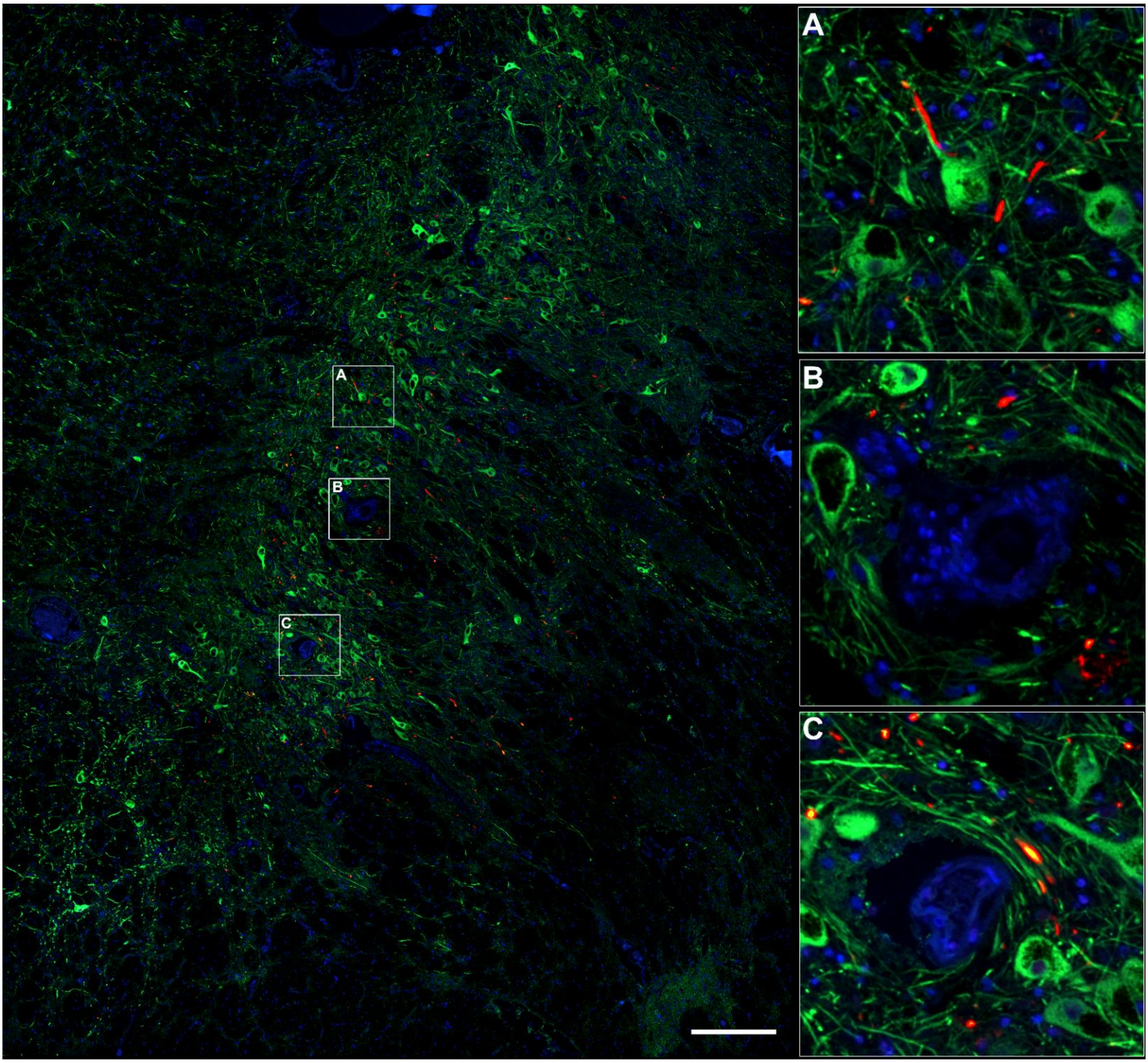

**Supplementary Figure 4.** Tyrosine Hydroxylase (green), Nucleocapsid Protein (red) and Hoechst (blue) immunofluorescent staining. Nucleocapsid protein immunoreactivity can be detected within the boundaries of the substantia nigra in the midbrain and may be found within tyrosine hydroxylase positive neurites and neuronal somata.
